# Innervation and Neuronal Control of the Mammalian Sinoatrial Node: A Comprehensive Atlas

**DOI:** 10.1101/2020.10.28.359109

**Authors:** Peter Hanna, Michael J. Dacey, Jaclyn Brennan, Alison Moss, Shaina Robbins, Sirisha Achanta, Natalia P. Biscola, Mohammed A. Swid, Pradeep S. Rajendran, Shumpei Mori, Joseph E. Hadaya, Elizabeth H. Smith, Stanley G. Peirce, Jin Chen, Leif A. Havton, Zixi (Jack) Cheng, Rajanikanth Vadigepalli, James Schwaber, Robert L. Lux, Igor Efimov, John D. Tompkins, Donald B. Hoover, Jeffrey L. Ardell, Kalyanam Shivkumar

## Abstract

Cardiac function is under exquisite intrinsic cardiac neural control. Neuroablative techniques to modulate control of cardiac function are currently being studied in patients, albeit with variable and sometimes deleterious results. Recognizing the major gaps in our understanding of cardiac neural control, we sought to evaluate neural regulation of impulse initiation in the sinoatrial node as an initial discovery step. Here, we report an in-depth, multi-scale structural and functional characterization of the innervation of the sinoatrial node (SAN) by the right atrial ganglionated plexus (RAGP) in porcine and human hearts. Combining intersectional strategies including tissue clearing, immunohistochemical and ultrastructural techniques, we have delineated a comprehensive neuroanatomic atlas of the RAGP-SAN complex. The RAGP shows significant phenotypic diversity of neurons while maintaining predominant cholinergic innervation. Cellular and tissue-level electrophysiologic mapping and ablation studies demonstrate interconnected ganglia with synaptic convergence within the RAGP to modulate SAN automaticity, atrioventricular (AV) conduction and left ventricular (LV) contractility. For the first time, we demonstrate that intrinsic cardiac neurons influence the pacemaking site in the heart. This provides an experimental demonstration of a discrete neuronal population controlling a specific geographic region of the heart (SAN) that can serve as a framework for further exploration of other parts of the intrinsic cardiac nervous system (ICNS) in mammalian hearts and for developing targeted therapies.

## Introduction

The nervous system intricately regulates almost every aspect of cardiac physiology and profoundly influences pathophysiological adaptions to disease (1). Cardiac autonomic control is achieved by a nuanced interplay between the extra-cardiac neural circuits that interface with the intrinsic cardiac nervous system (ICNS), the clusters of ganglia found in ganglionated plexuses (GPs) in epicardial fat pads on the surface of the heart. These GPs contain afferent and efferent neurons that transduce information from and to the myocardium as well as local circuit neurons (2, 3). The ICNS coordinates cardiac reflexes in concert with inputs from the hierarchy of the cardiac neuraxis composed of cortical centers, brainstem, spinal cord and intrathoracic sympathetic ganglia and may serve as a nexus point for therapies for cardiovascular disease (3).

Understanding the control of the ICNS on discrete cardiac regions has implicated specific GPs with certain aspects of cardiac functions. For example, a GP in the intercaval region at the dorsal aspect of the right atrium (RA), known as the right atrial GP (RAGP), has been shown to mediate vagal influences over the sinoatrial node (SAN) (4–7). Following the description of the human ICNS, a series of comparative anatomic studies show that the RAGP is conserved among mammalian species with fibers originating from the epicardial fat pad and coursing toward the SAN (7). It is now recognized that GPs do not merely serve as relay stations housing postganglionic parasympathetic neurons, but as integration centers that are interconnected to affect cardiac function (8).

Clinically, there has been recent interest in neuroablative and neuromodulatory interventions at the level of the ICNS. For example, botulinum neurotoxin has been shown to reduce arrhythmia recurrence when injected into the ICNS in one study with 3-year follow-up (9). In contrast, the neuroablative approach evaluated in the Atrial Fibrillation Ablation and Autonomic Modulation via Thorascopic Surgery (AFACT) study did not demonstrate improved arrhythmia recurrence-free survival and, unfortunately, was associated with increased pacemaker implantation rates (10). Furthermore, while histologic studies in the past have been suggestive of neural pathology in the setting of cardiovascular disease such as sudden death (11), disruption of parasympathetic nerves has also been shown to increase ventricular arrhythmogenesis in a murine model (12). Given the eagerness to perform neuro-interventions for arrhythmias in the clinical setting with varying and sometimes deleterious effects, we set forth to perform foundational work to provide a guide to intrinsic cardiac neuroanatomy and physiological control.

Advancing our knowledge of the neural circuits controlling the cardiac impulse origin at the SAN is a logical neural-visceral link to explore the function of the ICNS. Recent advances in characterizing peripheral neural circuits will aid in elucidating the intrinsic cardiac neural control of cardiac function, and here we apply these techniques to understanding the interactions between RAGP and SAN (13). Because rodent cardiac ganglia do not appear as organized into discrete GPs as in higher order mammals and the study of large animal models is advantageous in translational arrhythmia research, we chose the Yucatán minipig as a large animal model (14–17). In this study, we employ a multidisciplinary approach to develop in-depth phenotypic and functional assessments of the ultrastructural to macroscopic neuroanatomic connections between the RAGP and SAN in porcine and human tissues.

## Methods

### Animals

Animal experiments were performed in accordance with the UCLA Institutional Animal Care and Use Committee, and euthanasia protocols conform to the National Institutes of Health’s Guide for the Care and Use of Laboratory Animals (2011). Data for all experiments were collected from 41 male and female Yucatán minipigs (> 3 months old).

### Human Cardiac Specimens

Use of a fresh frozen cadaveric heart (Donated Body Program, University of California Los Angeles) from a 97-year-old woman with history of coronary artery disease who died of cardiopulmonary failure was approved by the UCLA Institutional Review Board.

Three de-identified donor human hearts were approved for research by the Institutional Review Board (Office of Human Research) at the George Washington University (GWU). Human hearts were procured by Washington Regional Transplant Community and arrested in ice-cold cardioplegic solution in the operating room prior to delivery to GWU. An RAGP from a 31-year-old man with no cardiac history who died of anoxia (hanging) was immersion fixed in 4% paraformaldehyde (PFA) and used for histologic study. The specimen used for tissue clearing was from a 71-year-old Asian woman with history of diabetes mellitus and hypertension who died following a large intracranial hemorrhagic stroke. The heart used for optical mapping was from a 52-year-old Caucasian woman with history of hypertension and hyperlipidemia who died of stroke.

### Perfusion Fixation of Human Heart

A de-identified female donor heart without underlying cardiac disease underwent perfusion fixation with 4% PFA for tissue clearing. Three of the pulmonary veins and the inferior vena cava were sutured closed with one pulmonary vein and the superior vena cava serving as outflow of perfusate. The aortic root and pulmonary artery were cannulated with 24 Fr cannulas. Perfusion of 4% PFA via these cannulas was performed at flow rates to achieve approximately 80mmHg in the left ventricle and 20mmHg in the right ventricle. Perfusion was maintained at 4°C for 24 hours. The cannulas and outflow were then clamped, and the tissue was immersion fixed with agitation at 4°C overnight. The heart was then washed with 0.01M PBS perfusate for 30 minutes via cannulas followed by three 0.01M PBS washes for 30 min. The heart was then stored in 1x PBS/0.02% sodium azide at 4°C until processing for modified immunolabeling-enabled three-Dimensional Imaging of Solvent-Cleared Organs (iDISCO+) tissue clearing.

### Coronary Artery Contrast Injection and Imaging

The right side of the fresh frozen cadaveric human female cardiac specimen was used to determine the vascular supply of the RAGP. The right coronary artery was selectively cannulated. BriteVu media (Scarlet Imaging, Utah) was prepared by heating diH_2_O to 40-45°C. 700µL of BriteVu enhancer was added to 105mL of warmed water. After 1 minute, 35g of BriteVu (40% wt/vol) was added to solution and heated to 70-80°C for 10 minutes with agitation. The solution was allowed to cool to 45°C and pressure perfused using a peristaltic pump (PERIPRO-4HS, World Precision Instruments) via the right coronary artery at 50mL/min. The specimen was then immersion fixed in 4% PFA. Specimens were imaged using a Nikon D850 DSLR camera with Nikon AF Micro-NIKKOR 200mm f/4D IF-ED lens and scanned with a custom-built microCT at 50kV and 200μA with 260m resolution at the California Nanosystems Institute.

### Neural Tracing Experiments

For initial surgery, following sedation (induction: ketamine (10mg kKg-^1^ IM)/midazolam (1mg Kg-^1^ IM), maintenance: isoflurane 1-2% inhalation) and intubation, a right unilateral thoracotomy was performed by dividing the pectoral muscle, making a small incision in the pericardium, and exposing the right atrial-superior vena cava junction. Then 5mg of an analog of 1,1’-dioctadecyl-3,3,3’,3’-tetramethylindocarbocyanine perchlorate (*FAST* DiI, Invitrogen, D7756) in 250µL of 100% methanol (MeOH) was injected using a 27-guage needle into the SAN region. A chest tube was placed, and the incision was closed. Immediately prior to removal, the chest tube was aspirated. Tissues were harvested in a terminal procedure at least 3 weeks later as described below.

### Porcine Tissue collection

Following sedation (induction: tiletamine-zolazepam 6mg Kg^−1^ IM, maintenance: isoflurane 1-2% inhalation) and intubation, we performed a midline sternotomy and exposed the heart. A heparin bolus of 5000U IV was administered, and the pig was then placed in ventricular fibrillation with application of a 9V battery to the surface of the heart. The heart was explanted and syringe-flushed with heparinized normal saline (5U mL^−1^) via the transected aorta. The area of interest (RAGP-SAN region) was then excised and rinsed in heparinized saline. Whole-mount preparations of RAGPs for electron microscopy (EM) analysis were isolated under a dissection microscope and immersion fixed overnight at 4°C in a 2% PFA and 2.5% glutaraldehyde solution in 0.12M Millonigs buffer (MB). RAGPs for histologic, tissue clearing and immunohistochemical studies were immersed in 4% PFA at 4°C with agitation overnight. The following day, the tissues were washed three times for 30 minutes each in 0.01M PBS and stored in either 0.01M PBS and 0.02% sodium azide solution at 4°C for histologic and tissue clearing studies or 0.01M PBS with 20% sucrose and 0.02% sodium azide for at least 1 week for cryopreservation for immunohistochemical studies. SAN region for RNAscope (ACD) was embedded in Optical Cutting Temperature compound (OCT, TissueTek) in a cryomold and placed on dry ice. Tissue was stored at −80°C until processed. RAGPs for gene expression profiling were separated from the SANs, immersed in 0.01M PBS at room temperature (RT) for 30 seconds and transferred to 25% OCT, followed by 50% OCT and then embedded in 100% OCT and placed in a cryomold. The cryomold was placed in a MeOH dry ice bath for flash freezing.

### Histologic Studies

Porcine and human epicardial RAGP fat pads and SANs were paraffin embedded and serially sectioned at 5µm thickness. We obtained representative histologic sections and stained epicardial RAGP fat pad and SANs with hematoxylin and eosin (H&E) and Masson’s trichrome. Slides were imaged on Aperio AT Turbo scanning machine at 20x.

### iDISCO+ Tissue Clearing

Portions of porcine and human cardiac tissue were stained and cleared using the iDISCO+ protocol as previously described (13, 18). Fixed tissue underwent graded MeOH dehydration (20, 40, 60 and 80% MeOH in diH_2_O [vol/vol]) for 1 hour each at RT with agitation, washed twice with 100% MeOH for 1 hour at RT and incubated in 66% dichloromethane/33% MeOH overnight at RT with agitation. The following day, tissues were washed twice in 100% MeOH for 1 hour at RT, chilled at 4°C and then bleached in 5% H_2_O_2_ in MeOH (vol/vol) overnight at 4°C. Then, the tissues were rehydrated with a graded MeOH series and washed in 0.01M PBS for 1 hour each at RT with agitation. The tissues were washed in 0.01M PBS with 0.2% Triton X-100 for 1 hour at RT. Tissues were then stained by permeabilizing in 0.01M PBS, 0.2% Triton X-100, 20% dimethyl sulfoxide (DMSO) and 0.3M glycine and blocking in 0.01M PBS with 0.2% Triton X-100, 10% DMSO and 5% donkey serum, each for 2 days at 37°C with agitation. Tissues were then incubated in primary antibodies rabbit anti-PGP9.5 (Abcam, ab108986, 1:200) diluted in 0.01M PBS with 0.2% Tween-20 and 10mg ml^−1^ heparin (PTwH) for 1 week at 37°C with agitation. Hearts were washed 4 to 5 times in PTwH overnight at room temperature. Secondary antibodies donkey anti-rabbit Cy3 (Jackson ImmunoResearch, 711-165-152,1:300) for 1 week at 37°C with agitation. Primary and secondary antibodies were replenished midway through staining. Hearts were again washed in PTwH 4-5 times overnight at RT. To clear, tissues were dehydrated with a graded MeOH series as above and incubated in 66% dichloromethane/33% MeOH for 3 hours at room temperature with agitation. Tissues washed twice in 100% dichloromethane for 15 minutes at room temperature and cleared, stored, and imaged in benzyl ether (Millipore Sigma, Catalog 108014; refractive index: 1.55).

### Immunohistochemical Methods for Sections

Fixed specimens for porcine hearts containing the RAGP fat pad, SAN and right atrium were shipped to ETSU at 4°C in PBS, 20% sucrose and 0.02% sodium azide solution. The fat pad and SAN with adjoining RA were dissected separately, frozen on dry ice, and stored at −80°C until sectioning. 30µm thick sections were cut at −20 to −25°C using a Leica CM3050S cryostat (Leica Microsystems Inc.) and collected on charged slides. Tissues were sectioned in a plane parallel to the epicardial surface, and sections were collected in a sequence that yielded 8 sets of sections that each spanned the entire thickness of the specimen. Each set of tissue sections was stored at −20°C until further processing.

Slide-mounted tissue sections were immunostained at RT for markers using standard methods of fluorescence immunohistochemistry, as described previously (19). Briefly, slides were rinsed with PBS (pH 7.3), incubated for 10 minutes in PBS containing 0.4% Triton X-100 and 0.5% bovine serum albumin (BSA), and blocked for 2 hours in PBS containing 1% BSA, 0.4% Triton X-100, and 10% normal donkey serum (NDS; Jackson ImmunoResearch Laboratories). Each section was then incubated in blocking buffer containing one or more of the primary antibodies listed in **Table S1**. Sections were washed again and incubated in blocking buffer before application of secondary antibodies. Species-specific donkey secondary antibodies conjugated to Alexa Fluor 488, 555, 594 or 647 (Jackson ImmunoResearch Laboratories) were applied at a 1:200 dilution in PBS containing 0.4% Triton X-100 and 1% BSA, and sections were incubated for 2 hours before final washing in PBS. After final washes with PBS, cover glasses were applied with Citifluor (Ted Pella, Inc.) or SlowFade Gold antifade reagent (Life Technologies Corporation). Specific staining did not occur in negative control sections processed without the addition of the primary antibodies.

Staining for tyrosine hydroxylase (TH) was amplified using a biotin-streptavidin method. Biotin-SP conjugated donkey anti-goat secondary antibody (Jackson ImmunoResearch Laboratories) was applied in place of regular secondary antibody and incubated for 2 hours. Sections were then washed with PBS and incubated for 2 hours in streptavidin conjugated Cy3 (Jackson ImmunoResearch Laboratories).

### RNAscope

The OCT-embedded SAN was equilibrated in the Leica cryostat at-20°C for 1 hour, and 20µm sections were mounted on charged slides to undergo RNAscope per the manufacturer’s protocol. Briefly, sections were dried at −20°C for 2 hours and stored at −80°C until used. Slides were then immersed in chilled 4% PFA for 90 minutes at 4°C followed by 2 rinses with 0.01M PBS. Sections were dehydrated in a graded series of 50%, 70%, 100% and 100% ethanol (EtOH in diH_2_O; vol/vol) for 5 minutes each. The slides were then allowed to air dry for 5 minutes at RT and baked at 60°C for 30 minutes. A barrier around the section was created using a hydrophobic barrier pen and allowed to dry over 5 minutes. Slides were loaded in a humidity control tray, and 5 drops of H_2_O_2_ were added followed by a 10-minute incubation at RT. Slides were then washed with diH_2_O twice. Five drops of Protease IV were added to each section and incubated for 30 minutes at RT in humidity control tray. Slides were then washed with 0.01M PBS twice. Five drops of each probe mix were added to each slide. Probe mixes included a negative control, positive control mix (*PPIB*, *POLR2A-C2* and *UBC-C3*) and *HCN4*. Slides were baked at 40°C for 2 hours in a humidity control tray. In this and the subsequent washing steps, slides were washed with wash buffer for 2 minutes twice. Five drops of Multiplex FL v2 Amp 1 were added to each slide and baked at 40°C for 30 minutes. After washing, 5 drops of Multiplex FL v2 Amp 2 were added to slides, and the slides were baked at 40°C for 30 minutes. Slides were and 5 drops of Multiplex FL v2 Amp 3 were added to each slide and baked for 15 min at 40°C. Slides were washed and then 5 drops of Multiplex FL v2 HRP-C1 were added to each slide and baked for 15 minutes at 40°C. After washing, 150µL of Opal 690 dye (1:1500 in DMSO) was added to each slide and incubated for 30 minutes at 40°C. Following another wash, 5 drops of Multiplex FL v2 HRP blocker were added, and slides baked at 40°C for 15 min. After washing, 5 drops of Multiplex FL v2 HRP-C2 were added to each slide and baked at 40°C for 15 minutes. After another wash, 150µL of Opal 570 dye to each slide and incubated for 30 minutes at 40°C. After washing, 5 drops of Multiplex FL v2 HRP blocker were added to each slide and incubated at 40°C for 15 minutes. Following washing, 4 drops of DAPI was added to each slide, incubated for 30 seconds at RT, and then tapped dry. One drop of ProLong Gold Antifade Mountant was added to each slide and coverslipped. Slides were left to dry overnight in the dark and stored at 4°C until imaging.

### Immunohistochemical Method for Ganglion Whole-Mount Preparations

After electrophysiologic recordings, isolated ganglia were fixed in 4% PFA overnight. Fixed tissue was washed in 0.01M PBS 3 times for 1 hour each and stored in 0.01M PBS with 0.02% sodium azide. The tissue was blocked in 0.01M PBS, 0.02% sodium azide, 0.2% Triton X-100 and horse serum for 4 hours at RT with agitation. The following primary antibodies were used: rabbit anti-PGP9.5 (1:500), sheep anti-TH (1:200) in 0.01 M PBS, 0.02% sodium azide and 0.2% Triton X-100 with agitation for 2 nights. Tissues were washed with a solution of 0.01M PBS with 0.02% sodium azide every hour for 3 hours. Tissue was incubated in secondary antibodies diluted in 0.01M PBS with 0.1% Triton X-100 and 0.02% sodium azide for 2 nights at room temperature with agitation. The following secondary antibodies were used: donkey anti-rabbit Cy3 (1:200), donkey anti-sheep 488 (Jackson ImmunoResearch, 713-545-147, 1:200). Secondary staining with streptavidin conjugated ATTO-647N was used to visualize neurobiotin filling (1:500). Stained tissue was rinsed in 0.01M PBS with 0.02% sodium azide every hour for 3 hours, mounted on glass slides, and coverslipped.

### Imaging and Image Processing

Slides were viewed under fluorescence illumination with an Olympus BX41 microscope equipped with an Olympus DP74 digital camera and cellSens software (Olympus America Inc., Center Valley, PA, RRID:SCR_016238). For quantitative studies of the RAGP, SAN and adjoining RA, slides were evaluated with a Leica TCS SP8 Confocal Microscope with 10x, 20x and 40x objective lenses (Leica Microsystems Inc.). Confocal images were collected at a resolution of 1024 × 1024 using 488, 552 and 638nm laser lines. Stacks spanned tissue thicknesses of 25-33µm unless otherwise noted. Figures were created using maximum intensity projection (MIP) images for individual channels and merged images. For quantitative studies of the neurochemical phenotype of neurons in the RAGP, confocal images were collected for every ganglion (≥ 3 neurons) in one set of section for each pig. For quantitative studies of nerve fiber density in the SAN and right atrium, ten 30x confocal images from separate fields, usually on different sections and slides, were collected for each pig.

Each iDISCO+-cleared tissue was placed in a chamber (SunJin Lab) filled with benzyl ether on a slide, and a coverslip applied. Tilescan and Z stack images were obtained using a Zeiss LSM 880 confocal laser scanning microscope with a Fluar 5x/0.25 M27 Plan-Apochromat and 10x lens. Images were obtained at a resolution of 1024 × 1024 using 488 and 561nm laser lines. Z-axis step size was commensurate with Nyquist sampling based on numerical aperture of the specified objective. Pinhole was set to 1 airy unit. Stitched images were analyzed in Zeiss Zen Black SR and Bitplane Imaris 9.5.1 for 3D visualization.

In situ hybridization sections and RAGP whole-mount preparations were imaged with an upright laser scanning confocal microscope (Zeiss AxioExaminer/LSM 880). For the RNAscope studies, 8-bit tile scan images were obtained using a Plan-apochromat 20x/0.8 objective lens. For the whole-mount preparations, tilescan and Z-stack images (8-bit) were acquired using both low-power (W Plan-Apochromat 10x/0.5 M27 75mm) and high-power (Plan-Apochromat 63x/1.40 Oil DIC M27) objectives to scan whole tissue (cm^2^) or individual ganglia (μm^2^), respectively. Z-axis step size and pinhole were set as mentioned above. Laser excitation of fluorophores (DAPI: 405nm diode laser; Opal 570 and Cy3: 561 nm diode pumped solid state laser; Alexa Fluor 488: 488nm argon laser; Opal 690 and ATTO 647: 633nm HeNe laser) was optimized for maximum gain without oversaturation of the detector. Muscle autofluorescence was obtained using the 488nm laser line. Stitched images were analyzed in Zeiss Zen Black SR.

### Image Analysis

The RAGP fat pads from pigs injected with DiI into the SAN region were sectioned as described above. Sections were first evaluated without attaching cover glasses, and images of every ganglion were collected using the Olympus fluorescence microscope and camera. This approach was taken because some DiI would be lost during subsequent immunohistochemical staining. After documenting DiI labeled neurons, the sections were immunolabeled for PGP9.5 to identify all neurons in the ganglia. For each ganglion, we were then able to count DiI-labeled neurons and the total number of neurons (i.e., PGP9.5-positive).

Quantification of RAGP neurons singly or doubly labeled for specific neuronal markers was done using Stereo Investigator software v2019.1.1 (MicroBrightField, Inc., RRID:SCR_002526). Unique markers were used to differentiate total neuron count, single and double labeled neurons.

Nerve density in cardiac regions was determined by using ImageJ software to evaluate single channel confocal images that were labeled for VAChT, TH and NPY. Nerve density was calculated as the area occupied by nerves as a percentage of the entire image area and reported as % area.

Colocalization of two markers in nerve fibers was evaluated using MIP confocal images. The colocalization rate was quantified in select images using a module of the Leica LASX Software (RRID:SCR_013673). This module displays a two-channel image as a scatter plot that is used to adjust thresholds so that only points of pixel overlap are retained, and the background is removed. Colocalization rate is reported as a percentage of colocalization area/total area.

### Transmission Electron Microscopy

After MB and double-distilled H_2_O (ddH_2_O) washes, the tissues were fixed in 1% osmium solution diluted in ddH_2_O followed by dehydration in ethanol and propylene oxide. The tissues were blocked and embedded in Epon plastic resin (Ted Pella). Semi-thin cross sections (0.5μm) were obtained and labeled with 1% toluidine blue solution diluted in ddH_2_O for overview visualization of RAGP organization at light microscopy level using a Nikon Eclipse E600 microscope and Nikon camera DS-Fi3. Ultrathin sections (70nm thickness) were collected on formvar coated grids (Ted Pella), counterstained with uranyl acetate and lead citrate, and examined under a Tecnai G2 Spirit Twin transmission electron microscope (FEI, ThermoFisher Scientific) operating at 80_KV and images acquired with a Gatan Orius SC 1000B digital camera (Gatan) for detailed characterization of the RAGP organization.

### Cryosectioning and Staining for Laser Capture Microdissection-High Throughput RT-PCR

RAGP was consecutively sectioned at 40µm thickness along the superior-inferior vena caval axis, imaged and mounted on PPS membrane slides (Leica Microsystems, Catalog 11600294). An average of 400 slides were generated per RAGP. Slides were fixed in ice-cold 100% ethanol for 1 minute followed by 4 minutes of staining with 0.0001% Cresyl Violet (ACROS Organics, AC229630050). Slides were dehydrated using 95% and 100% ethanol followed by xylene for 1 minute each. RNAse inhibitor (Invitrogen SUPERase-In RNase Inhibitor, Catalog AM2696) was added to all aqueous reagents to preserve RNA quality.

### Laser Capture Microdissection

Samples were collected immediately post staining using laser capture microdissection (LCM; Arcturus, ThermoFisher). Single neurons were visualized in the 0.0001% cresyl-violet stained sections using a brightfield and triple filter. Samples were collected on LCM caps and stored at −80℃. Samples were lysed at the time of HT-qRTPCR experiments with lysis reagents as described below.

### Mapping LCM samples onto the reconstructed 3D stack

2D blockface images generated for each tissue section during cryosectioning were used to generate a 3D volume in Tissue Mapper software (MicroBrightField, Inc.). Images taken during LCM sample collection were referenced to corresponding sections in the 3D stack. Anatomical features and neurons were assigned markers enabling extraction of specific XYZ coordinates. Blockface images and acquisition images can be found at DOI: 10.26275/56h4-ypua.

### High-throughput real-time PCR

Single RAGP neurons in lysis buffer (Cells Direct Lysis Buffer, Invitrogen) were processed for reverse transcription using SuperScript VILO Master Mix (Thermo Fisher Scientific) followed by real-time PCR for targeted amplification and detection using the Evagreen intercalated dye-based approach to detect the PCR-amplified product. Intron-spanning PCR primers were designed for every assay using Primer3 (20) and BLAST (21). Genes selected spanned a variety of neuronal functions, signal transduction and cell-type identification. Per the standard BioMark protocol, cDNA samples were processed for 22 cycles of specific target amplification of 283 genes using TaqMan PreAmp Master Mix as per the manufacturer’s protocol (Applied Biosystems). Real-time PCR reactions were performed using 96.96 BioMark Dynamic Arrays (Fluidigm) enabling quantitative measurement of multiple mRNAs and samples under identical reaction conditions. Each run consisted of 30 amplification cycles (15 seconds at 95°C, 5 seconds at 70°C, 60 seconds at 60°C). Real-Time PCR Analysis Software (Fluidigm) was used to calculate Ct values. Twenty-one 96 × 96 BioMark Arrays were used to measure gene expression across the 422 (before QC) single-cell samples. The same serial dilution sample set was included in each chip to assess batch effects. Samples from each RAGP were run across 3 chips to obtain data on 283 genes per sample. Each set of chip runs for a given animal contained overlapping assays that served as technical replicates to evaluate chip-to-chip variability, which was minimal.

### High-throughput-qPCR data analysis

Melt-curve analysis was used to assess quality of qRT-PCR results. Samples with >30% failed reactions and genes with >20% failed reactions were excluded from analysis. An additional 10 samples were outliers and excluded from analysis. A total of 405 single-cell samples and 246 different gene assays were included in the final analysis. Raw Ct values for individual samples were normalized against a median expression level of a subset of 140 robustly expressed genes (genes with greater than 60% working reactions) across all animals to obtain −∆Ct values. The vector of median sample expression value was chosen over potential reference genes based on comparison of stable expression across all samples against known housekeeping genes using the ‘selectHKs’ function in the NormqPCR package in R. The −∆Ct data were then rescaled using the median across all samples within a gene using the following equation:

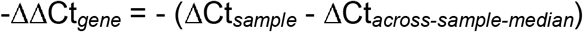

The raw and normalized dataset is available online as a Gene Expression Omnibus dataset (GEO reference ID: GSE149212). Pearson correlation was performed to compare gene expression levels and generate heatmaps.

### Intracellular Recordings from Intrinsic Cardiac Neurons

Tissue was isolated as above and placed in ice-cold physiologic salt solution (PSS) containing in mM: 121 NaCl, 5.9 KCl, 1.2 NaH_2_PO_4_, 1.2 MgCl_2_, 25 NaHCO_3_, 2 CaCl_2_, 8 D-glucose; pH 7.4 maintained by 95% O_2_-5% CO_2_ aeration. Ganglia of the RAGP were isolated from the surface of the inferior vena cava-right atrium junction in proximity (~5-10mm) to the SAN. Isolated whole ganglia were pinned to the SylGard (Dow Corning) floor of a glass bottom petri dish. RAGP neurons were observed with an upright microscope (Zeiss AxioExaminer) equipped with a 5x and a 40x water-immersion objective.

Intracellular measurements of membrane potential were obtained following technique described previously (22). Preparations of ganglia were continuously superfused (6–7 ml/min) with PSS maintained at 32–35°C with a thermostatically controlled heater. Neurons were impaled with borosilicate-glass microelectrodes filled with 2M KCl + 2% Neurobiotin (80-160MΩ; Vector Labs). Membrane voltage was recorded with a Multiclamp 700B amplifier connected to a Digidata 1550B data acquisition system. pCLAMP 10 software (Molecular Devices) was used for data acquisition and analysis. Depolarizing and hyperpolarizing currents were applied through the recording electrode to characterize neuronal membrane properties. Hyperpolarizing steps (−100pA, 500ms) were given to measure steady state input resistance. Depolarizing current steps (0.1–0.5nA, ∆ 100pA, 500ms duration) were used to assess neuronal excitability. Inclusion criteria for analysis included a resting membrane potential less than −40mV, a holding current of greater than −100pA, and cells must generate an action potential.

Concentric bipolar stimulation electrodes (FHC) were placed onto interganglionic nerves. Graded stimulus shocks (100µs) in 50-100μA steps, from 0 to 800μA were used to generate stimulus recruitment curves (Master 8 and IsoFlex optical Isolation unit, AMPI). Five to 20 stimuli were delivered at each stimulus intensity, with an interval of 3 seconds between stimuli. At least two stimulating electrodes were used to activate discrete nerve bundles into a single ganglion.

### Ablation of Porcine RAGP

Surgery was performed as above except a clamshell thoracotomy was performed for improved exposure of the RAGP. Following thoracotomy, anesthesia was changed to α-chloralose (50 mg/Kg intravenous bolus administration followed by continuous infusion at 10 mg/Kg/hr IV). The left (LCV) and right cervical vagi (RCV) were dissected free in the neck and encircled with platinum nerve cuff electrodes. A quadripolar His catheter (Abbott) was placed via the right external jugular vein and advanced until a His signal was visualized while a quadripolar catheter was inserted via the right femoral vein and advanced to the right atrium. The left (LSG) and right stellate ganglia (RSG) were dissected free and either platinum needle electrodes or cuff electrodes were placed. All stimulations were performed individually using a Grass S88 Stimulator via PSIU6 constant current isolation units. Square wave stimulation pulses (10Hz frequency, 1ms duration, 0.1-15mA) were delivered individually to each vagus nerve, and stimulus threshold was defined as the stimulation current strength that was sufficient to elicit a 10% decrease in heart rate. Current amplitude was increased to 3 times threshold for all subsequent vagal nerve stimulation (VNS). Square wave stimulation pulses (4ms duration, 4Hz frequency, 0.1-15 mA) were delivered individually to each ganglion or sympathetic chain. Stimulus threshold was defined as the stimulation current strength that was sufficient to elicit a 10% increase of left ventricular end-systolic pressure (LVESP) or heart rate. Current amplitude was increased to three times threshold for all subsequent sympathetic stimulations. Right atrial pacing was performed using a Micropace EPS320 system at 2 times capture threshold at a rate at least 20% greater than the resting heart rate during the last 10 seconds of each VNS.

Continuous 12-lead surface ECG and intracardiac ECG data were recorded using a Prucka CardioLab (GE Healthcare). For ARI recordings, a custom 56-electrode sock, placed over both ventricles, was connected to the Prucka CardioLab system to identify regional activation recovery intervals (ARI). Global ventricular ARIs were calculated via customized software ScalDyn M (University of Utah), as described previously (23). ARI, a surrogate marker of local action potential duration, was calculated as the difference between the first minimal dV/dt in the QRS complex to the first maximal dV/dt of the T wave.

Systolic LV pressures were assessed by using a 5F pigtail, 12-pole conductance-pressure catheter connected to an MPVS Ultra processor (Millar Instruments, Inc.) placed in the left ventricle via carotid or femoral artery sheath under ultrasound guidance. Pressure was continuously monitored and recorded throughout experiments. Systolic LV function was assessed by LV end-systolic pressure (LVESP) and the maximum rate of LV pressure change (dP/dt_max_). Data were acquired using the Cambridge Electronic Design System and Spike2 software. Heart rates and atrial-His (AH) intervals were measured in Spike2 software.

After baseline stimulations at 3 times threshold of the bilateral stellate ganglia and cervical vagi were performed, the RAGP was ablated using a Stryker Sonopet ultrasonic aspirator. After the RAGP was ablated, repeat VNS with atrial pacing and SGS were performed.

### Electrical Recordings of SAN Region

After thoracotomy, instrumentation of bilateral vagi and stellate ganglia and placement of right atrial and His catheters as above, a custom-made 64-channel multielectrode array (Neuronexus) was positioned over the sulcus terminalis and electrical recordings were acquired using an Alpha Omega system. The bilateral vagi and stellate ganglia were individual stimulated at three times threshold as above and recordings at the SAN performed. Then, platinum needle electrodes were inserted to the RAGP and stimulated using the Grass S88 Stimulator via PSIU6 constant current isolation units. Square wave stimulation pulses (50Hz frequency, 0.1ms duration, 0.1-15mA) were delivered until a heart rate decrease of at least 10% was noted. Analysis of SAN recordings was performed using ScalDyn M software. Activation times (ATs) were measured from the beginning of the P wave to the first minimal dV/d*t* in the atrial electrogram.

### Right Atrial Wedge Preparation and Optical Mapping

For optical mapping of the human SAN, the entire RA was carefully isolated and the right coronary artery was cannulated with a custom flexible plastic cannula. Bleeding vessels were tied off, and the tissue was stretched across a custom frame and transferred to a vertical bath of warmed (37°C) and oxygenated (95% O_2_/5% CO_2_) Tyrode’s solution (in mM: 128.2 NaCl, 4.7 KCl, 1.05 MgCl_2_, 1.3 CaCl_2_, 1.19 NaH_2_PO_4_, 20 NaHCO_3_, 11.1 Glucose). Adequate perfusion was maintained for the duration of the experiment for tissue viability, and sensing electrodes were pinned into the bath for pseudo ECG recordings using LabChart. The preparation was electromechanically uncoupled with blebbistatin (10-15µM) and fluorescently stained with di-4-ANEPPS. Optical action potentials (OAPs) were captured with MiCAM05 CMOS cameras with high spatial and temporal resolutions (100_×_100 pixels, 1kHz sampling frequency; SciMedia). During recordings, the endocardial surface of the tissue was illuminated with a 520 nm LED (Prizmatix), and emitted fluorescence was captured through a 650 nm long-pass filter (Thorlabs).

RHYTHM, a custom-made MATLAB program designed for optical mapping data analysis, was used for the creation of optical activation maps (OAPs) and identification of the leading pacemaker sites. OAPs were spatially (5 × 5 pixel neighborhood) and temporally (low pass Butterworth filter at 150–200_Hz) filtered, and 60 Hz noise was removed. Baseline fluorescent drift was also removed with a first- or second-order fitted curve, and activation maps were created by identifying 50% of the maximum OAP amplitude. Stimulation of the RAGP was performed with a bipolar stimulus electrode placed on the epicardial fat pad at frequencies of 50 Hz and 100 Hz with a pulse width of 200µs.

### Statistics

GraphPad Prism 8.4.2 was used for data analysis and graph generation. Based on preliminary GP ablation experiments, we estimated we would need 4 pigs to detect a 40% bradycardia with a standard deviation of 20% at an α level of 0.05 at 80% power in paired difference analysis for the functional experiments. Sample sizes are included in figure legends or text. Data are presented as mean ± SD or as median (interquartile range [IQR]) for non-normally distributed data. Comparisons for all pairs were conducted using a two-tailed paired Student’s t-test or Wilcoxon matched-pairs signed rank test for non-normally distributed data. P <0.05 was considered statistically significant.

### Data Availability

All data supporting findings in this study are available in the SPARC Data Portal and from the corresponding authors upon reasonable request. Raw and processed HT-qPCR data of single neurons from the pig RAGP have been deposited in the GEO database under accession code: GSE149212. All sample acquisition images, raw and processed transcriptomic data, and annotations pertaining to 3D spatial location are publicly available in the sparc.science repository with the identifiers 10.26275/56h4-ypua, 10.26275/kabb-mkvu, 10.26275/5jki-b4er, 10.26275/qkzi-b1mq, and 10.26275/255m-00nj.

## Results

### Neuroanatomy and vascular supply of the RAGP

The RAGP is located at the dorsal aspect of the intercaval region and lateral to the right pulmonary vein complex in both pig and human (**Fig. 1**). A discrete epicardial fat pad containing the RAGP could be reliably found in porcine and donor human cardiac tissues, and histologic examination illustrated that ganglia are found suspended within adipose tissue as well as close to the fat-RA muscle interface (**Fig. 1B-C, G-H and Movie S1**). To visualize the neural tissues in three dimensions, we applied a tissue clearing technique, iDISCO+, to generate optically transparent tissue for large volume imaging (13, 18, 24). iDISCO+-clearing of the porcine and human RAGP and immunostaining with pan-neuronal marker protein gene product 9.5 (PGP9.5) demonstrates that numerous ganglia, found across the thickness of the RAGP fat pad, are interconnected by nerve fiber bundles (**Fig. 1D, I-K and Movies S2-4**). We injected the SANs of two pigs with neural tracer, *FAST* DiI, to assess for direct projections from the RAGP to the SAN (**Fig. 1A**). When evaluated after three weeks for tracer transport, we observed robust labeling of neurons and nerve fibers in the RAGP and, in many cases, several neuronal processes were evident on individual neurons (**Fig. 1E**). Three weeks was deemed sufficient as labeling of neurons within the right stellate ganglion (RSG) was identifiable (**Fig. S1**). The extent of DiI labeling was consistent between pigs, with 21.5% of PGP9.5-positive neurons labeled in one animal and 24% labeled in the other. These neurons were distributed in ganglia across the depth of the RAGP with an enrichment of SAN-projecting neurons at the aspect of the RAGP closest to the SAN (**Fig. 1F**). Injection of contrast media into the right coronary artery of a human cadaveric specimen followed by microCT imaging confirmed that the vascular supply of the RAGP is via the sinus nodal branch of the right coronary artery (**Fig. 1L and Movie S5**).

**Fig. 1.**
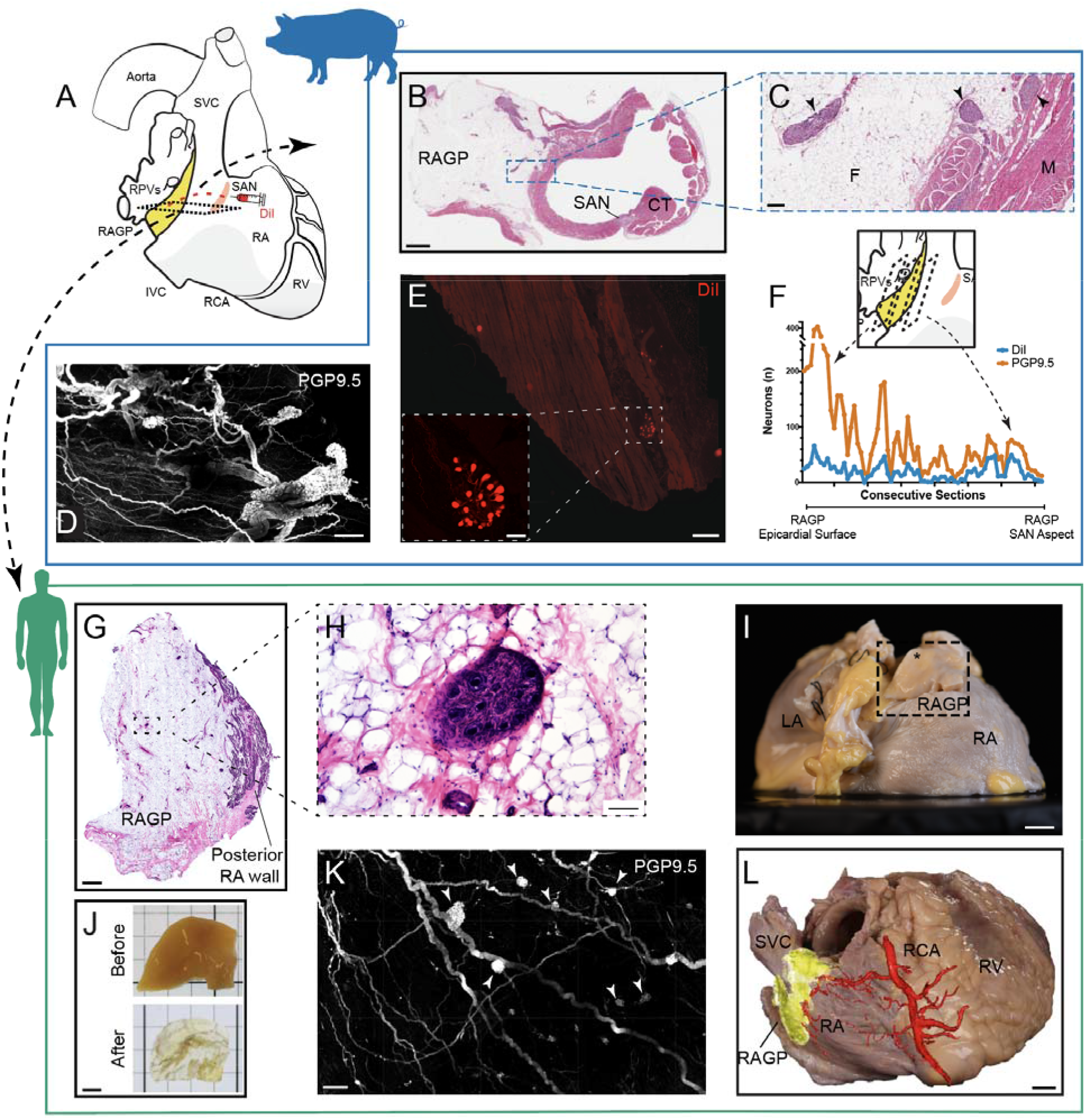
Neuroanatomy and vascular supply of the porcine and human right atrial ganglionated plexus (RAGP) in relation to the sinoatrial node (SAN). **A** Schematic of human heart demonstrating proximity of RAGP to SAN. DiI was injected into the porcine SAN followed by tissue harvest of RAGP 3 weeks later. **B** Hematoxylin and eosin (H&E) staining of porcine RAGP-SAN region in a male pig imaged at 20x. CT: crista terminalis. **C** Ganglia (arrowheads) suspended in adipose tissue as well as at fat-right atrial muscle interface at higher magnification. F: fat, M: muscle. **D** Maximum intensity projection (MIP) image of iDISCO+-cleared RAGP from a male pig illustrating cluster of interconnected ganglia with PGP9.5 (white). **E** Example of DiI-labeled neurons and nerve fibers in a ganglion in the RAGP. Higher magnification of DiI-labeled ganglion is shown (inset). **F** Quantification of DiI- and PGP9.5-positive and total neurons from a female pig. Eight representative sets of sections were collected from each RAGP in this study, and this data was obtained from one set that spanned the entire RAGP (1.8cm). **G** H&E staining of human RAGP fat pad at the posterior RA wall. **H** Ganglion suspended in fat at higher magnification. **I** Right posterior oblique view of gross anatomy of a second human RAGP dissected (inset) for tissue clearing. Dissection contained epicardial fat (*) with adjacent muscle. LA: left atrium. **J** Portion of human RAGP before and after iDISCO+ tissue clearing. **K** MIP of iDISCO+-cleared portion of human RAGP demonstrating clusters of interconnected ganglia (arrowheads) with PGP 9.5 (white). **L** Photograph of right posterior oblique view of a third human heart with overlay of microCT image of contrast-media-injected right coronary artery showing RAGP is supplied by the sinoatrial nodal artery. Scale bars are 2mm (**B, G**), 200µm (**C**), 500µm (**D, K, E**), 100µm (**E inset, H**), 1cm (**I, L**), 5mm (**J**).

### Macroscopic and cellular organization of the porcine RAGP

Most of the ganglia are in the adipose tissue near the atrial muscle and between superficial muscle layers (**Fig. 2A-D**). The typical fat pad has surface dimensions of approximately 2 × 1.5cm and penetrates to a depth of about 1.5 to 2cm. Ganglia are present throughout the depth of the fat pad and vary in size, containing from 3 to almost ~170 neurons with substantial variation in size of neurons with cross-sectional area ranging from 599.0 to 3766µm^2^ (**Fig. 2E-H**). Single representative sets of sections from the RAGP fat pad contained from around 750 to 1,500 neurons, and the estimated total number of neurons per RAGP ranged from 6,000 to 12,000.

**Fig. 2.**
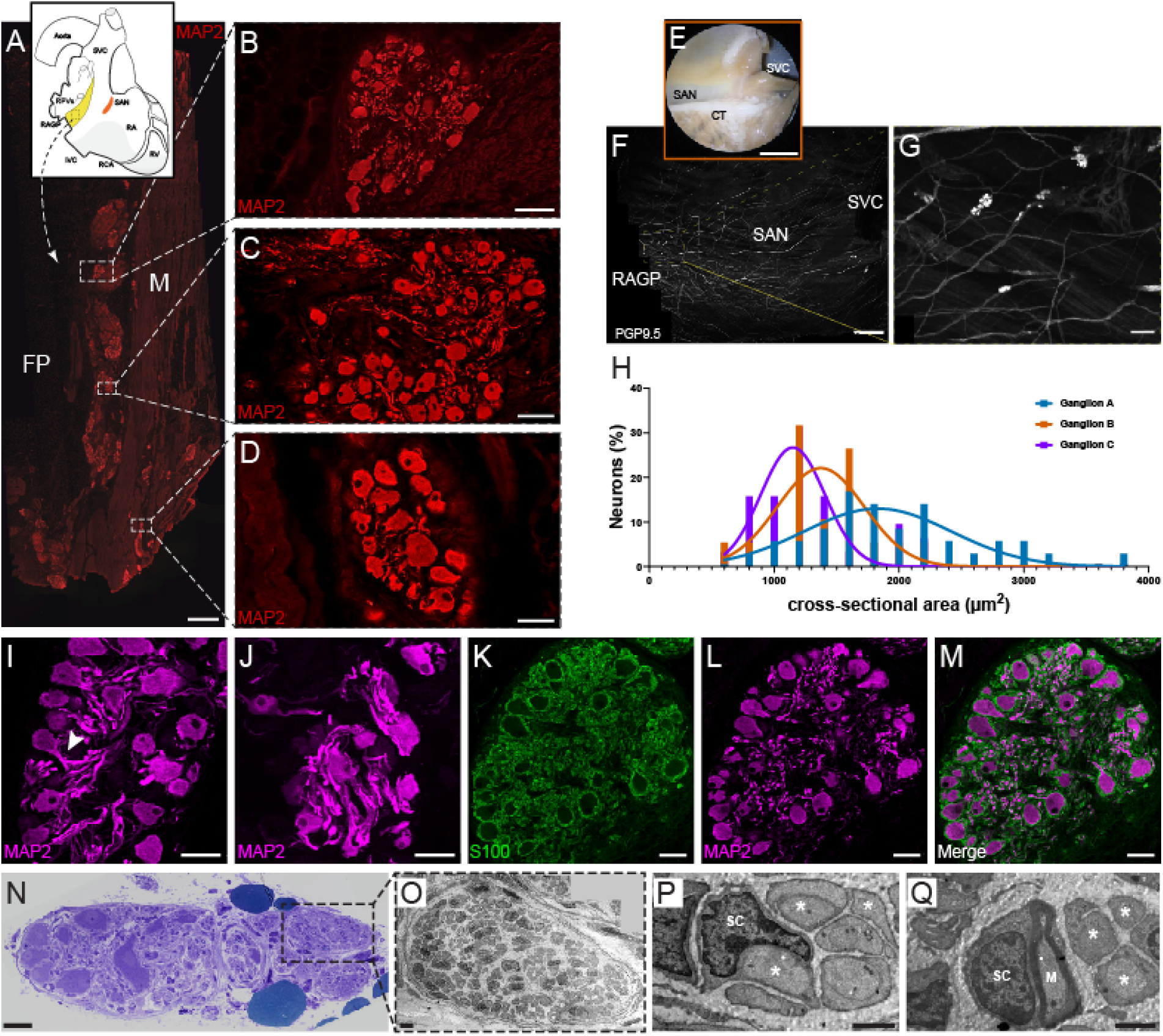
Immunohistochemical and electron microscopic characterization of distribution and morphology of ganglia in the RAGP fat pad. **A** Photomontage of section that was immunostained for the pan-neuronal marker microtubule-associated protein 2 (MAP2; red). This figure was created from 165 overlapping 10x fluorescence images. **B-D** Three ganglia shown at higher magnification. Most ganglia occur near the interface between the fat pad (FP) and atrial muscle (M), although some occur between muscle bundles (**D**). **E** Photograph of SAN region of RAGP-SAN tissue from a male pig that underwent whole-mount staining (**F**). **G** Examples of three ganglia identified in **F** that underwent quantification of cross-section area (**H**). **I-J** Ganglia stained for MAP2 contain many multipolar neurons, some with one or more thick processes and a variable number of thinner processes. Some of the latter appear short, likely due the plane of sectioning and others are longer. Occasionally, axons bifurcate after leaving the cell body (**I**, arrowhead). **K-M** Neurons in the porcine RAGP are surrounded by satellite glial cells. Ganglia were double labeled for the glial marker S100 and the pan-neuronal marker MAP2 (**K-L**), and corresponding overlay images (**M**) show that S100-positive cells and processes surround MAP2-positive perikarya, axons and other processes. **N** Light microscopy image of toluidine blue staining of RAGP shows several distinct sub-compartments, some with ganglionic neurons and some without neurons. **O-Q** Ultrastructural detail of RAGP fascicle with numerous unmyelinated fibers (*) and some myelinated fibers. Note Schwann cell nuclei in **P**(SC) and one to one close relationship between myelinated axon (M) and Schwann cell in **Q**. Five porcine RAGPs (3 male, 2 female) were evaluated in the immunohistochemical studies and 4 (2 male, 2 female) in the electron microscopy studies. Scale bars are 1mm (**A**), 50µm (**B, D, I-M**), 100µm (**C**), 1cm (**E**), 2.5mm (**F**), 200µm (**G**), 30µm (N) and 0.5µm (**O-Q**).

In contrast to ganglia of smaller species like the mouse, cardiac ganglia of the pig contained primarily multipolar neurons (**Fig. 2I-J**). Each neuron is surrounded by satellite glial cells that also envelop intraganglionic nerve processes (**Fig. 2K-M**). Additionally, some ganglia and other sites in the fat pads contained clusters of small intensely fluorescent (SIF) cells (**Fig. S2**). Transmission electron microscopy studies of porcine RAGP demonstrated an organizational structure where the RAGP is divided into several distinct compartments and separated by fibrous tissues (**Fig. 2N-Q**). Ganglionic neurons were present in small clusters that also included satellite cells as well as myelinated and unmyelinated fibers.

### Neurons in the porcine RAGP and synaptic inputs to the ganglia are primarily cholinergic

We evaluated whether neurons in the porcine RAGP exhibit a cholinergic phenotype and receive extensive cholinergic input from presumed preganglionic vagal efferent nerve fibers. This was confirmed using a well-validated antibody to the vesicular acetylcholine (ACh) transporter (VAChT; **Table S1**), a specific cholinergic marker. The antibody labeled about 99% of the neural cell bodies in the RAGP and revealed intense, punctate staining of presumed preganglionic nerve varicosities in the ganglionic neuropil (**Fig. 3A-C**). Double staining for VAChT and pan-neuronal marker microtubule-associated protein 2 (MAP2) showed that many of these varicosities are found near neuronal cell bodies and their processes.

**Fig. 3.**
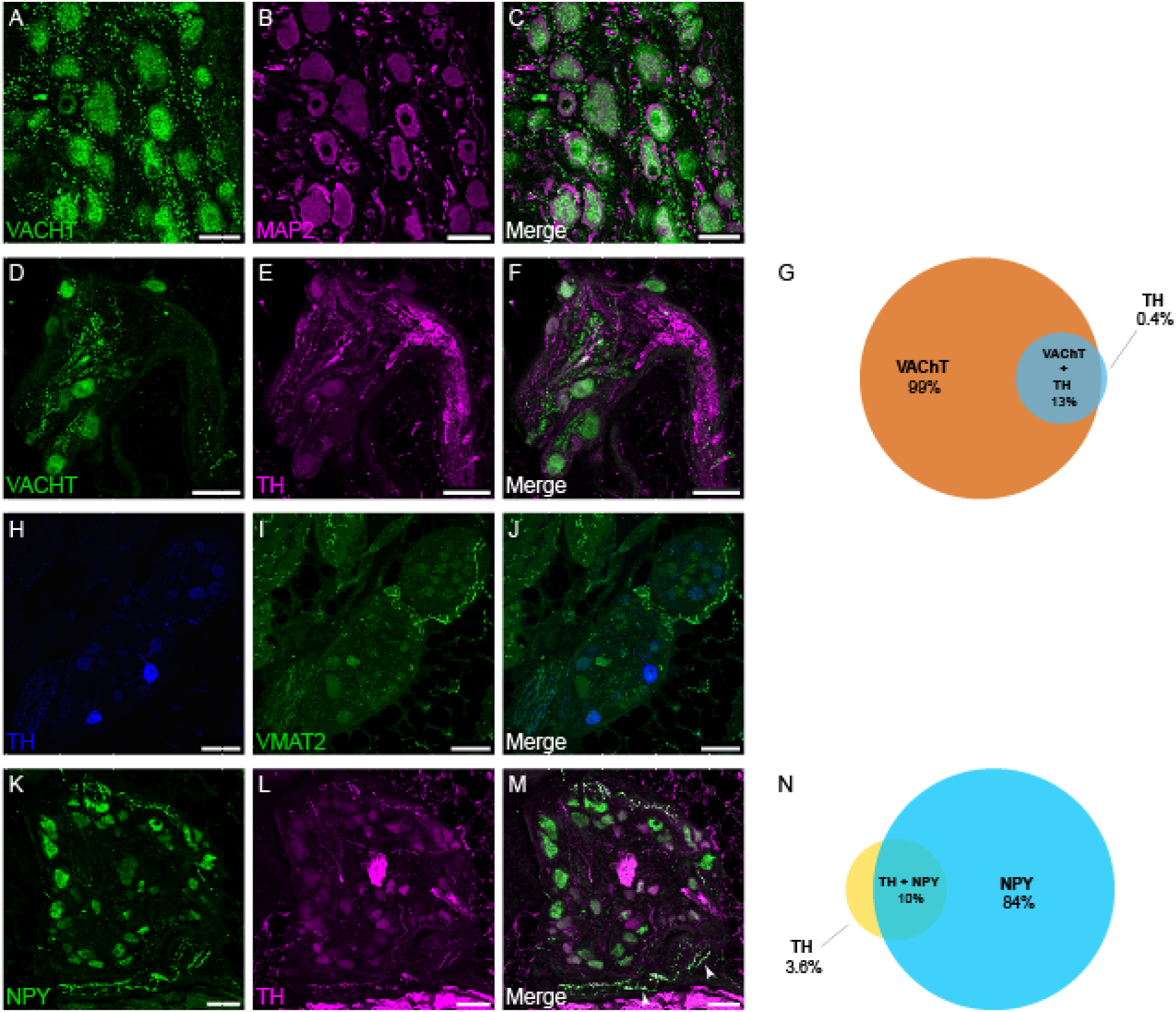
Ganglia in the RAGP contain cholinergic neurons and receive cholinergic input. Confocal images of ganglia double labeled for the cholinergic marker VAChT (**A**) and for MAP2 (**B**). VAChT appears as granular staining of neuronal cell bodies and varicose staining in the neuropil. Neuronal cell bodies and processes are more fully delineated by MAP2 staining. **C** Overlay images show colocalization of VAChT and MAP2 in cell bodies and the close association of VAChT-positive varicosities to cell bodies and nerve processes. **D-F** Confocal images of ganglia double labeled for VAChT (**D**) and for noradrenergic marker TH (**E**). Merged images indicate some neurons that contain VAChT and TH (**F**). Venn diagram illustrates proportion of VAChT-positive, TH-positive and both VAChT- and TH-positive neurons (**G**). A few RAGP neurons stain for TH (**H**) but usually lack the essential noradrenergic marker VMAT2 (**I**), although positive staining for VMAT2 does occur in nerve fibers. **J** Merged image shows that TH-positive cell bodies lack staining for VMAT2. Confocal images of ganglia double labeled for NPY (**K**) and TH (**L**) illustrate that many neurons show moderate to intense, granular labeling for NPY, but the same neurons lack TH. In contrast, NPY and TH colocalized in some nerve. **M** Merged image shows that nerve fibers labeled for NPY and TH (arrowheads) are sparse within the ganglia and not in close apposition to ganglionic neurons. Venn diagram illustrates proportion of NPY-positive, TH-positive and both NPY- and TH-positive neurons (**N**). Each value is based on analysis of confocal images from at least 8 microscopic fields in porcine RAGP (n = 4; 2 male, 2 female). Scale bars are 50µm (**A-C**), 100µm (**D-F, H-M**).

### Some neurons in porcine RAGP express non-cholinergic neurotransmitter markers

We evaluated neurochemical coding of the porcine RAGP (n = 4; 2 male, 2 female) in a series of double and triple labeling experiments. Staining for VAChT and TH showed that about 13% of the cholinergic neurons also contain the noradrenergic marker TH (**Fig. 3D-G**), consistent with prior studies in other animal models (25), and a small percentage of neurons expressed TH without VAChT. However, most TH-positive neurons lacked staining for vesicular monoamine transporter 2 (VMAT2; **Fig. 3H-J**), which is essential for the storage and release of norepinephrine. This contrasts with noradrenergic neurons of the stellate ganglion, which stain intensely for both TH and VMAT2 (**Fig. S3A-C**). Nevertheless, a single ganglion with noradrenergic neurons was identified in the RAGP from one out of four pigs evaluated. This ganglion was innervated by VAChT-positive cholinergic nerve fibers and contained typical noradrenergic neurons that stained for TH, VMAT2 and NPY (**Fig. S3D-L**). Double labeling of RAGP for TH and NPY showed that 84% of ganglionic neurons showed granular staining for NPY (**Fig. 3K-N**). While TH-positive nerve fibers were abundant in nerve fiber bundles, only scattered nerve fibers within the ganglia stained for TH or NPY. Such fibers appeared to pass through or ramify within the ganglia, occasionally passing close to neurons (**Fig. 3M**). However, neither TH nor NPY was localized to varicose nerve fibers that surrounded RAGP neurons.

We also evaluated expression of neuronal nitric oxide synthase (nNOS), somatostatin and VIP. While we did not detect any neurons expressing somatostatin or VIP, nNOS was expressed in 3% of the neurons evaluated (**Fig S4A-E**). No nerve fibers staining for nNOS were detected within ganglia or nerve bundles of the RAGP.

### Neurons of porcine RAGP are not surrounded by sensory nerve fibers containing CGRP and SP

Innervation of intrinsic cardiac neurons by putative nociceptive sensory nerve fibers that contain CGRP and SP has been identified in a wide range of species, including humans (19). Frequently, varicose nerve fibers staining for these neuropeptides surround the intrinsic cardiac neurons, providing prominent innervation (19). Although porcine RAGP contains many nerve fibers that stain for CGRP and SP, most of these occur in nerve bundles that connect to the ganglia. Relatively few CGRP- or SP-positive nerve fibers enter the ganglia, and only a few of these single fibers pass close to intrinsic cardiac neurons (Fig. S4F-I).

### Cholinergic innervation is more abundant than noradrenergic innervation in the SAN but not in RA myocardium

We examined the innervation pattern of the SAN and adjacent RA myocardium. iDISCO+-cleared porcine SAN stained with PGP9.5 showed dense innervation of the SAN with several ganglia identified in the SAN region as well as larger fibers along the longitudinal axis of the crista terminalis (**Fig. 4A-B and Movie S6**). The SAN region identified by anatomic landmarks was confirmed using serial sections and 3,3’-Diaminobenzidine (DAB) staining with immunostaining for pacemaker channel HCN4 (**Fig. 4C**) and for the periphery of the SAN region with connexin 43, a marker of working myocardium (**Fig. 4D**). Dense cholinergic innervation was noted in this area (**Fig. 4E**). HCN4 staining did not work well with immunofluorescence (**Fig. S5A-B**), but we were able to delineate the SAN region by assessing for HCN4 transcripts using RNAscope (**Fig. 4F-H**). Staining for VAChT and TH showed that cholinergic nerves were more abundant than noradrenergic nerves in the porcine SAN (**Fig. 4, I-K**), but both nerve types had comparable density in atrial myocardium (Figure **4L-N**). Quantitative evaluation of nerve density in confocal images from these regions established that cholinergic nerve density is about five-fold higher than noradrenergic nerve density in the SAN and is significantly less in contractile atrial muscle (**Fig. 4O**). Noradrenergic nerve density was more uniform between regions. Since some neurons in the RAGP contained VAChT and TH, we wanted to determine if TH also occurred in cholinergic nerves in the SAN or RA. Colocalization analysis showed that this was not the case. However, double labeling for TH and NPY showed that these markers were often colocalized in the SAN (**Fig. 4P-R**) and adjacent RA (**Fig. 4T-V**). Quantitative evaluation of staining for TH and NPY showed that the density of nerve fibers for each marker did not differ in SAN, and NPY-positive fibers were slightly more abundant that TH-positive fibers in RA, although this was not statistically significant (**Fig. 4S**). These markers were highly colocalized in both regions (**Fig. 4W**). Nerves containing NOS, VIP or CGRP/SP were not detected in the SAN or adjacent RA. Examination of blood vessels in stained sections of the SAN and RA demonstrated prominent perivascular innervation by cholinergic and noradrenergic nerve fibers (**Fig. S6A-D**). The noradrenergic nerves also contained NPY. Perivascular nerves lacked NOS and VIP but did contain CGRP and SP (**Fig. S6E-F)**.

**Fig. 4.**
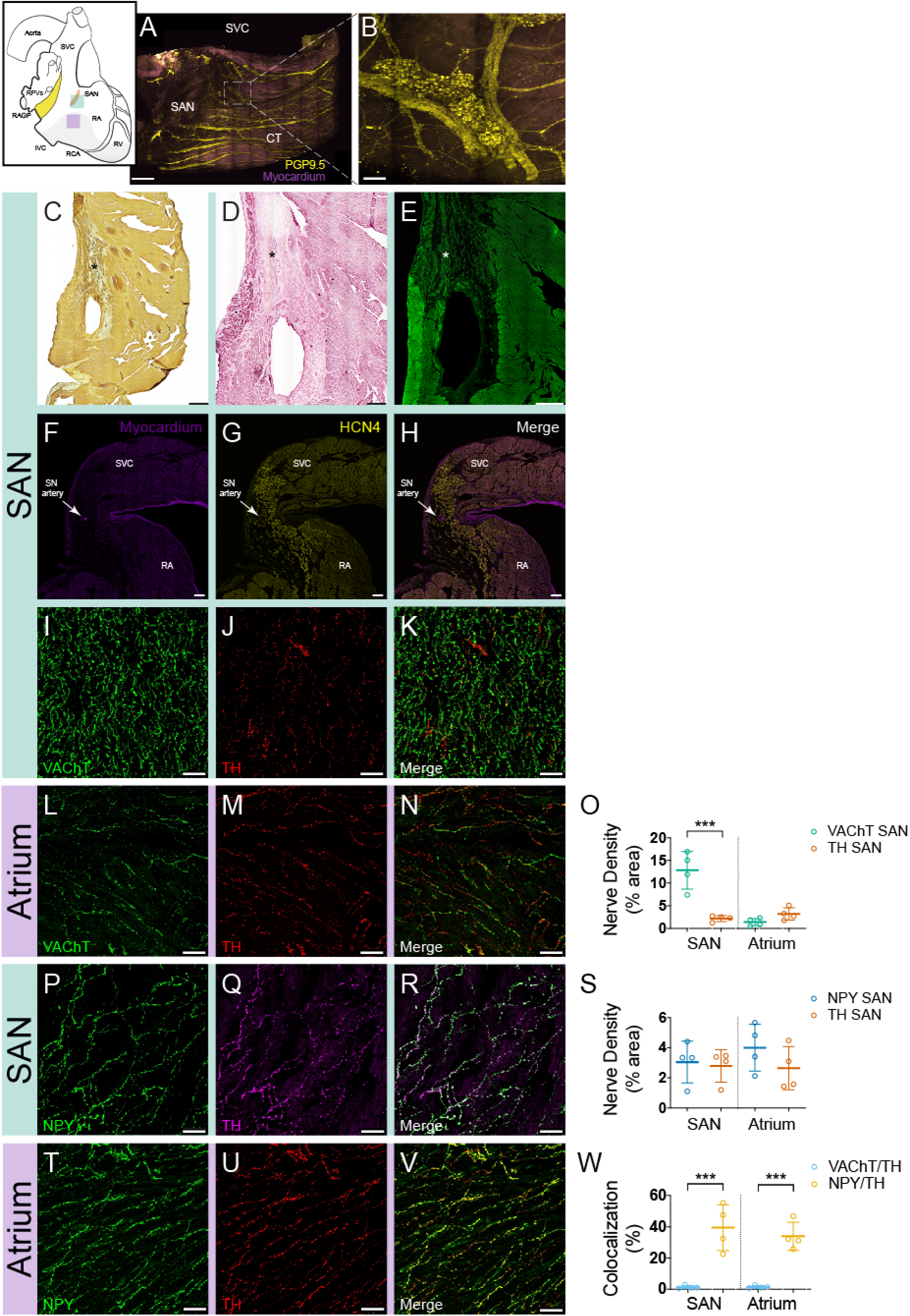
Cholinergic and noradrenergic nerves supply the SAN and atrial myocardium at different densities. **A** MIP of iDISCO+-cleared female porcine SAN illustrating ganglia and network of nerves (yellow). Purple: muscle autofluorescence. **B** Ganglia found in SAN region at higher magnification. **C** Sections of SAN (*) from a female pig that underwent DAB and HCN4 (brown puncta) staining demonstrates staining in the anatomic SAN region. H&E and connexin 43 staining of the working atrial myocardium border the region of the SAN region (*, **D**), which also demonstrates dense VAChT staining (**E**). RNAscope further confirmed *HCN4* gene expression (yellow) in the SAN region in a male pig (**F-H**). Sections of SAN (**I-K**) and right atrial myocardium (**L-N**) were double labeled for VAChT and TH. Confocal images from these sections show that cholinergic nerves (**I**) are more abundant than noradrenergic nerves (**J**) in the SAN, and that both nerve types have similar density in atrial myocardium (**L-N**). Merged images (**K, N**) show that VAChT and TH are not colocalized. **O** Density of cholinergic and noradrenergic nerves in the SAN and right atrial myocardium. Sections of SAN (**P-R**) and right atrial myocardium (**T-V**) were double labeled for NPY and TH. Confocal images from these sections show that staining for NPY (**P, T**) and TH (**Q, U**) labeled the same population of nerves in both regions. Merged images (**R, V**) show that NPY and TH have extensive colocalization. **S** Density of NPY-positive and TH-positive nerves in the SAN and right atrial myocardium. Sections were double labeled for these markers. **W** TH is not colocalized with VAChT but is colocalized with NPY in the SAN and right atrium. Each value is based on analysis of confocal images from at least 5 microscopic fields from 4 RAGPs (2M/2F). Scale bars are 2mm (**A, C-E**), 200µm (**B, F-H**) and 50µm (**I-N, P-R, T-V**). *** P≤0.001.

### Diverse adrenergic and cholinergic receptor gene expression in RAGP

To supplement the above neurochemical coding, we performed stochastic sampling of RAGP neurons for gene expression profiling. 405 neurons across 4 pigs (2 male, 2 female), mainly close to the posterior RA wall, were sampled and underwent high throughput PCR to evaluate the compilation of neurotransmitter receptors and ion channels expressed (**Fig. 5A, B**). Pan-neuronal markers were used as positive controls for the RAGP neurons (**Table S2**), which demonstrated a heterogeneity in the peptidergic, adrenergic and cholinergic, both nicotinic and muscarinic, receptor profiles (**Fig. 5C**). In addition to the diverse expression of sodium, calcium and potassium channel genes, genes encoding the HCN2 channels, typically found in nociceptive neurons, and HCN4 channels, which mediate *I*_*H*_ in neurons and *I*_*f*_ in the pacemaker cells of the heart, are expressed in RAGP neurons (**Fig. 5C-G**). While SP and CGRP was not identified by immunohistochemical methods in RAGP neurons, sensory marker SP (*TAC1*) and its receptor (*TACR1*) are highly expressed in HCN2-positive cells (**Fig. 5D-E**). Interestingly, in addition to pan-neuronal markers, neurotransmitters, and nicotinic receptors, *HCN2* gene expression was correlated with that of *HCN4*, which may suggest heteromerization of these proteins to form channels (**Fig. 5F-G**).

**Fig. 5.**
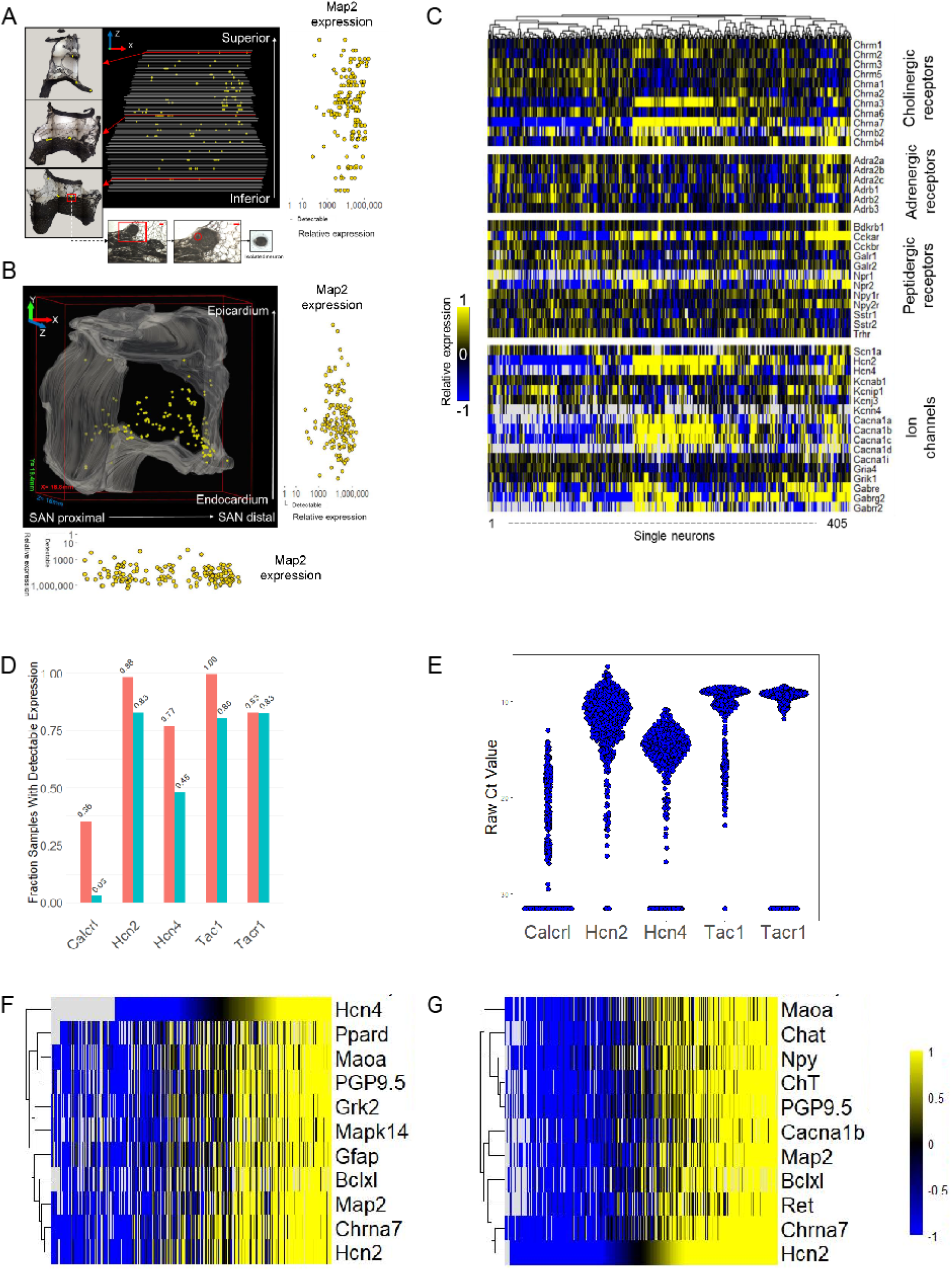
Diverse gene expression profile of porcine RAGP neurons. Projections of neurons isolated within the RAGP and their relative expression of pan-neuronal marker MAP2 in the XZ (**A**) and XY (**B**) planes. **C** Heat map demonstrating diverse profile of neurotransmitters, adrenergic, cholinergic and peptidergic receptors and ion channels in RAGP neurons. **D** A significant proportion of RAGP neurons exhibit high gene expression of HCN2 and HCN4. High expression of *TAC1* and *TACR1* genes, which are sensory markers, was identified. A smaller proportion expressed the receptor for CGRP (*Calcrl).* The range of expression of these genes is shown in panel **E**. The top 10 genes with correlated expression to HCN2 (Pearson correlation coefficient > 0.82; **F**) and HCN4 (Pearson correlation coefficient > 0.67; **G**) are shown. Note that high co-expression of pan-neuronal markers, neurotransmitters, nicotinic cholinergic receptor and downstream signaling molecules such as G Protein-Coupled Receptor Kinase 2 (*GRK2*).

### Ultrastructural imaging and cellular electrophysiological recordings demonstrate synaptic convergence and nicotinic transmission

In contrast to rodent intrinsic cardiac neurons, porcine RAGP neurons lack synaptic inputs to their somata (**Fig. 6A-B**). Instead, synaptic contacts are present on dendritic arbors (**Fig. 6C-E**). The synaptic boutons show primarily round clear synaptic vesicles with the presence of occasional dense core vesicles. To characterize these synaptic inputs, we performed cellular electrophysiologic measurements. We recorded from a total of 29 neurons isolated from 10 animals (5 male, 5 female). Mean resting membrane potential was −58 ± 7mV with input resistance of 95 ± 54 MΩ. Most cells (93%, 27/29) were phasic in nature, firing only a single action potential in response to intracellular injection of depolarizing current (100-500pA), while 2 cells fired 2-4 action potentials.

**Fig. 6.**
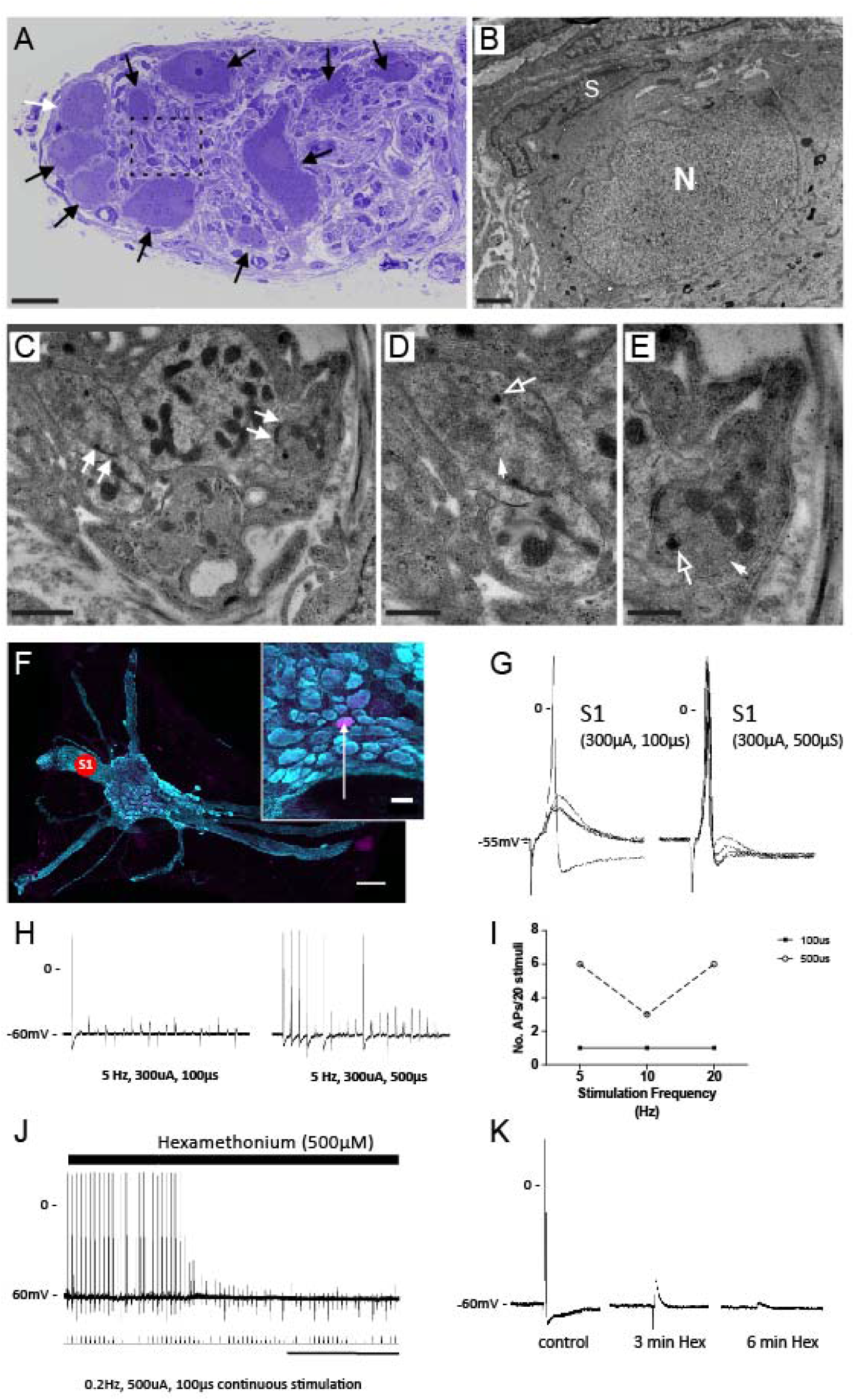
Convergence of cholinergic synaptic input at the RAGP. **A** Light microscopy image shows toluidine blue labeled RAGP neurons (arrows). **B** TEM image of neuron highlighted by white arrow in A. Note cell body nucleus (N) and satellite cell (S). No synaptic boutons are detected surrounding the cell body. **C-E** TEM of synaptic boutons in contact with dendritic profiles in the porcine RAGP (detected in regions highlighted by dashed line **A**). Note synaptic specializations in the form of active zones (arrows in **C**). The synaptic boutons contain clear rounded vesicles (closed arrows) and a few dense-core vesicles (open arrows). Each synaptic bouton-dendrite complex is in close apposition with surrounding satellite glial processes. **F** Representative ganglion within the RAGP. Note the many nerve bundles emanating from the ganglion. Concentric bipolar stimulating electrodes were placed on interganglionic nerves (S1) to elicit synaptic responses. Inset shows location of neuron (arrow) labeled with neurobiotin from the recording electrode. **G** Representative membrane potential responses recorded from the cell in F to stimulation at ‘S1’ are shown. There were no failures. Increasing stimulus current duration from 100 to 500 μS increased the probability of generating an action potential due to the larger evoked EPSP, demonstrating a convergence of inputs. **H** Synaptic efficacy was evaluated during continuous trains of presynaptic stimulation at the two pulse widths. Note the increased number of action potentials with the 500 μS pulse indicating fiber recruitment. **I** Synaptic efficacy (number of action potentials/stimulus frequency) was greater after increasing stimulus current duration. **J-K** The ganglion nicotinic antagonist hexamethonium blocked synaptically-mediated action potentials demonstrating a convergence of cholinergic inputs at RAGP neurons. Scale bar: (**A**) 30µm, (**B, C**) 1µm, (**D, E**) 0.5µm, (**F**) 250µm, (**F, inset**) 50µm, (**J**) 2 min.

Stimulation of interganglionic nerves produced both subthreshold and suprathreshold excitatory post-synaptic potentials (EPSPs). In 16 ganglion preparations, EPSPs were elicited by stimulation of two discrete nerve trunks demonstrating a convergence of inputs onto single cells. In 8 preparations, increasing stimulus intensity or duration at a single site recruited multiple inputs resulting in an increased amplitude of the EPSP and greater synaptic efficacy (**Fig. 6F-I**). In 5 preparations, treatment with the ganglion nicotinic antagonist hexamethonium (500μM) inhibited the nerve evoked potentials (**Fig. 6J-K**).

### Ablation of the RAGP mitigates VNS-induced bradycardia

To determine the control of the RAGP on cardiac function, the RAGP was ablated in 7 animals (4 males, 3 females; **Fig. 7A-B**). Following ablation, resting HR decreased from 100.5 ± 6.9bpm to 95.7 ± 7.5bpm (p = 0.031) but corrected sinus node recovery time (cSNRT) did not significantly change going from 106.9 ± 50.4ms to 105.0 ± 26.8ms (p = 0.921) (**Fig. 7C**). Ablation of the RAGP mitigated right cervical VNS (RCVNS)-induced bradycardia from 38.0 ± 8.2% to 5.9 ± 8.3% (p < 0.001) and left cervical VNS-(LCVNS-) induced bradycardia from 30.0 ± 5.5% to 5.6 ± 8.6% (p = 0.001) (**Fig. 7D, upper panel**). In addition to impacting sinoatrial rate, RAGP ablation abolished VNS-induced reductions in left ventricular (LV) contractility from 23.2 ± 5.5% to 5.2 ± 7.5% (p < 0.001) for RCVNS and from 31.7 ± 17.0% to 7.3 ± 5.4% for LCVNS (p = 0.014) (**Fig. 7D, lower panel**). RAGP ablation reduced right stellate ganglion stimulation (RSGS)-induced tachycardia from 32.2 ± 18.4% to 25.5 ± 14.2%, but this was not statistically significant (p = 0.066). Left SGS (LSGS) does not cause a significant tachycardia at baseline (3.4 ± 3.4%), and this was unchanged by RAGP ablation (2.3 ± 3.2%, p = 0.531). Similarly, RAGP ablation did not significantly impact SGS-induced increases in LV contractility (**Fig. 7D, lower panel**). RAGP ablation reduced RCVNS-induced prolongation of atrial-His (AH) interval, a measure of atrioventricular (AV) conduction, from 56.2 ± 38.5% to 6.7 ± 4.0% (**Fig. 7E**), during VNS with concurrent atrial pacing. In contrast, LCVNS-induced prolongation was not significantly affected, decreasing from 107.8 ± 33.1% to 66.0 ± 45.3% (p = 0.108). VNS-induced changes in ventricular activation-recovery interval (ARI), a surrogate marker of action potential duration, with concurrent atrial pacing did not significantly change following RAGP ablation (**Fig. S7**).

**Fig. 7.**
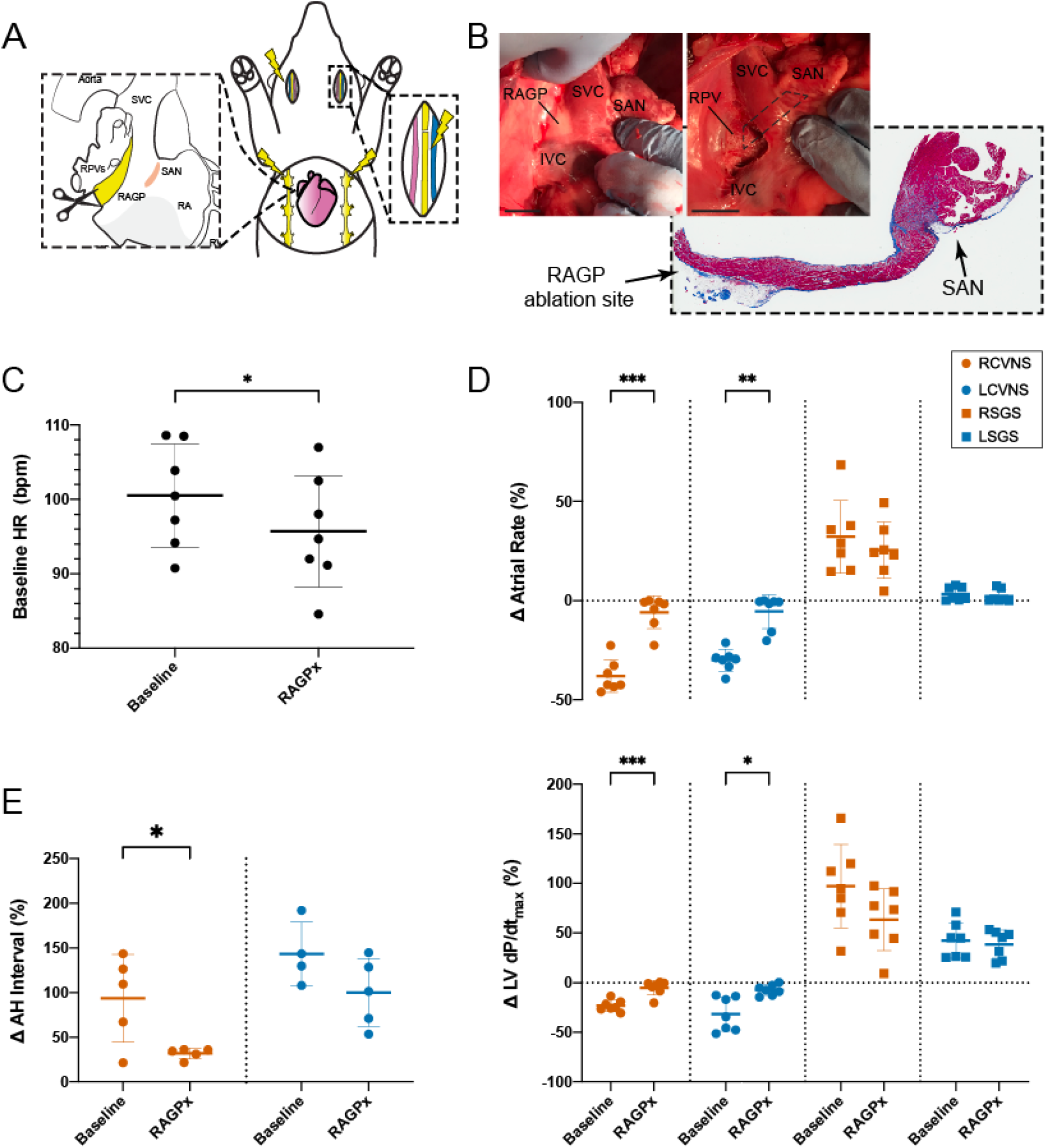
Ablation of the porcine RAGP mitigates VNS-induced bradycardia. **A** Schematic of RAGP ablation performed (n = 7) to assess impact of VNS- and SGS-induced changes on cardiac electrophysiology. **B** Gross photograph of porcine RAGP before and after ablation. Representative H&E section with trichrome staining of RAGP demonstrating destruction of epicardial fat pad without impacting SAN. RAGP ablation was followed by a reduced resting heart rate (**C**) and mitigated RCVNS- and LCVNS-induced effects on atrial rate and LV contractility as measured by maximal dP/dt (**D**) as well as AH interval (**E**). RSGS- and LSGS-induced changes in atrial rate and LV contractility were not statistically significant (**D**). Scale bars are 2cm (**B**). * P≤0.05, ** P≤0.01, *** P≤0.001.

### RAGP stimulation predominantly induces sinus bradycardia and pacemaker shift

At the macroscopic level, electrical stimulation of the RAGP was performed in a series of *in vivo* porcine experiments (n = 7; 6 male, 1 female). Stimulation of the RAGP was initially performed in pigs after bilateral cervical vagotomies, but it was generally difficult to elicit a response under these conditions. However, after complete decentralization with debranching from the vagosympathetic trunk, we were able to elicit at least a 10% bradycardia in 5 of 8 pigs from 88.7 (IQR 80.3-113.5 bpm) to 73.2 (IQR 57.3-100.0 bpm), including profound AV block (**Fig. 8A-C**). Additional effects of RAGP stimulation included initial bradycardia followed by tachycardia with up titration of frequency during stimulation, atrial fibrillation (AF) with slow ventricular response and increase in LV pressure with minimal HR increase (**Fig. S8A-B**). In addition to AF, electrical noise during high frequency stimulation precluded analysis of AH prolongation on intracardiac electrograms. Atrial electrograms were recorded using multielectrode arrays, but because of the large contribution of surrounding contractile myocardium to the electrogram, we were unable to ascertain shifts in pacemaker site (**Fig. S8C-E**). For this reason, *ex vivo* optical mapping of a human RAGP-SAN preparation was performed to assess nodal optical action potentials. Based on the vascular supply of the RAGP described above, a wedge preparation of the RAGP-SAN was maintained by coronary perfusion of the right coronary artery. Optical activation maps before and after RAGP stimulation, on tissue from a 52-year-old female human heart, showed an increase in HR from 49 to 113 bpm with 100Hz of stimulation and a superior shift in the pacemaker site (**Fig. 8D-F**).

**Fig. 8.**
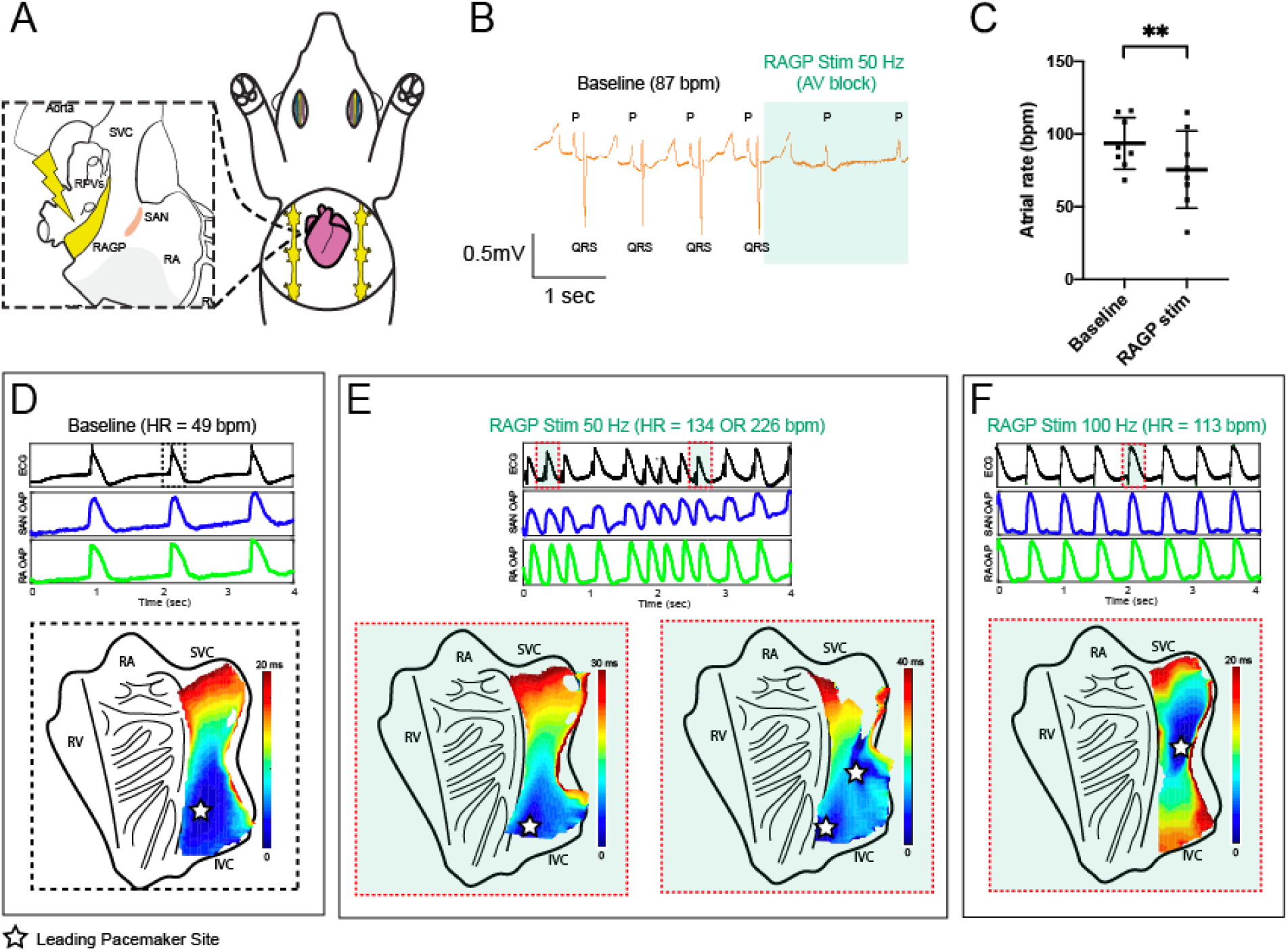
Stimulation of the porcine and human RAGP elicits SAN rate response with pacemaker shift. **A** Schematic of *in vivo* porcine RAGP stimulation with multielectrode array recordings in the SAN region. **B** Electrocardiogram demonstrating significant atrial rate reduction from 87 to 61 bpm and AV block during RAGP stimulation (50Hz, 0.1ms and 0.9mA). **C** RAGP stimulation (50Hz, 0.1ms) caused significant sinus bradycardia from a median of 88.7 (interquartile range [IQR] 80.3-113.5 bpm) to 73.2 (IQR 57.3-100.0bpm). Representative optical activation maps of an *ex vivo* sinoatrial nodal (SAN) preparation from a 52-year-old female human donor heart before (**D**) and during (**E, F**) stimulation of the RAGP. Intrinsic firing rate of the *ex vivo* SAN increased from 49 bpm to at least 134 bpm at 50Hz (**E**) and to 113 bpm with 100 Hz of stimulation (**F**). Stars indicate location of the leading pacemaker site on the endocardial side of the tissue. Corresponding ECG and optical action potential (OAP) recordings are shown below the activation maps. RA: right atria, RV: right ventricle, SVC: superior vena cava, IVC: inferior vena cava. ** P≤0.01.

## Discussion

We present a comprehensive assessment of the intrinsic cardiac neural control of the impulse origin in the mammalian heart. Using a multi-scale approach, we demonstrate 1) interconnected ganglia spanning the RAGP fat pad supply the SAN with an enrichment of neurons close to the SAN; 2) predominantly cholinergic innervation of the SAN; 3) heterogeneity in gene and protein expression profiling to identify putative neuronal subpopulations; and 3) the RAGP mediates primarily bilateral vagal inputs to exert profound functional control of the SAN as well as the AVN and LV (**Fig. 9**).

**Fig. 9.**
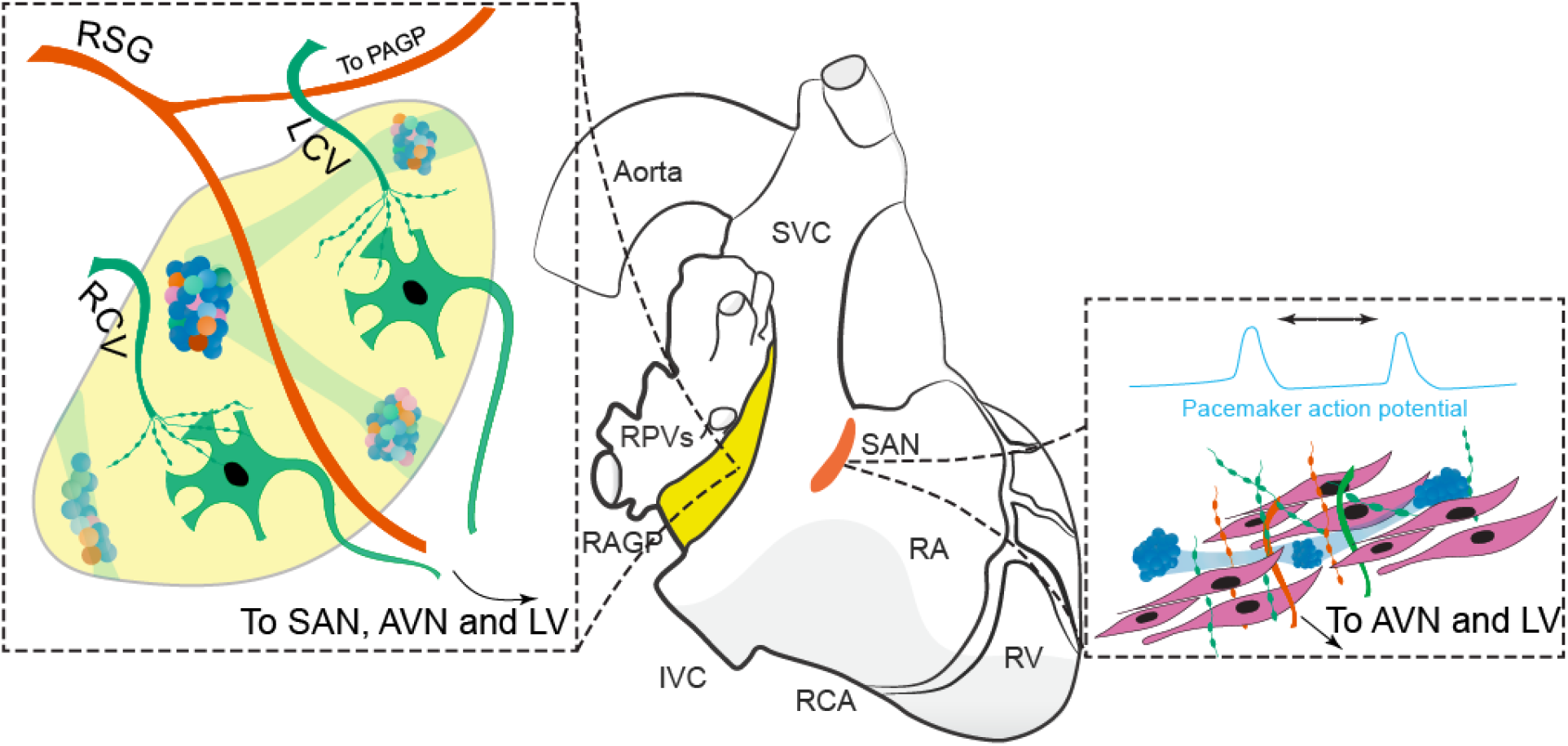
Intrinsic Cardiac Neural Control of the Mammalian Sinoatrial Node. Schematic diagram showing neuroanatomic connections between the RAGP and sinoatrial node and other cardiac regions as well as inputs from extrinsic nerves. Preganglionic neurons from the left (LCV) and right cervical vagi (RCV) synapse on postganglionic neurons within the RAGP. Postganglionic sympathetic fibers from the right stellate ganglion (RSG) pass through the RAGP but there are also fibers that likely course via the posterior atrial GP (PAGP). Note the phenotypic diversity of intrinsic cardiac neurons in the ganglia found in the RAGP. Ganglia are found near the SAN, and neural inputs to the SAN modify the sinoatrial rate and pacemaker site. Nerve fibers from the RAGP also supply the AVN and LV.

Within the paradigm of GPs acting as local control centers that serve specific cardiac regions, the RAGP has been postulated as exerting influence over the SAN. While prior studies have suggested RAGP neurons supply the SAN (26), the presence of SAN-projecting neurons throughout the RAGP was confirmed and quantified with the *in vivo* injection of a neural tracer at the porcine SAN. We were able to apply a tissue clearing technique to perform large volume imaging in porcine and human cardiac tissues to illustrate the dense neural network found in the RAGP fat pad with ganglia distributed throughout the adipose tissue as well as embedded in the posterior RA wall. This finding helps inform the epicardial versus endovascular approaches to targeting the GPs for arrhythmia treatment. We also identified distinct ganglia in the porcine SAN region, which has previously been variably described in the mouse, rat, guinea pig, cat, dog, and human, but whose function have yet to be elucidated (25, 27–30). Prior descriptions of catecholaminergic cells in ganglia near the SAN raise the possibility that these ganglia are involved in phenomena such as post-vagal tachycardia (31, 32) or the biphasic response of initial HR reduction followed by acceleration during ICNS stimulation (33). Large nerve fibers also tracked the crista terminalis, along which the impulse has been shown to preferentially propagate after exiting the SAN (34). In addition to characterizing neuronal populations, our immunohistochemical and ultrastructural studies demonstrate the close association of glial cells with RAGP neurons and intraganglionic nerve processes. As higher level of serum S100B following atrial fibrillation ablation has been associated with long-term treatment success, indicating that acute neural damage promoted freedom from arrhythmia recurrence (35), additional study of this cell population may prove critical in understanding ICNS control.

In the SAN region of multiple species, a dense meshwork of neurons has been characterized by immunohistochemical methods as predominantly cholinergic, although adrenergic-, nitrergic- and peptidergic-positive fibers have also been identified in the rabbit and human (19, 25, 28, 36, 37). Protein expression profiling indicates predominantly cholinergic neurons in the RAGP, but we describe the potential for expanded phenotypic diversity of RAGP neurons based on RNA analysis. For instance, neurons that expressed TH did not appear to be classically adrenergic, based on a lack of VMAT2 expression, and their function needs to be elucidated. Surprisingly, NPY, which is typically considered a cotransmitter for norepinephrine, was expressed in VAChT-positive cholinergic neurons in the RAGP and, conversely, in TH-positive fibers in the atrial myocardium. Peptidergic, putative sensory markers CGRP and SP were found in nerve fibers tracking through the ganglia but not the portion of the atrium we evaluated, which raises the question of how sensory information is transduced to RAGP neurons. Interestingly, transcripts of SP and its receptor are expressed within RAGP neurons but were not associated with significant protein expression on immunohistochemical study. While mechanical stimuli are thought to be transduced by the RAGP, the limbs of the Bainbridge reflex, the pathway by which atrial stretch induces HR increase, remains ill-defined (38, 39).

In addition to immunohistochemical study, prior work suggested the presence of adrenergic and cholinergic receptors in the GP based on response to neurochemicals; we are now able to provide a comprehensive library of genes expressed in intrinsic cardiac neurons (40, 41). Using high-throughput gene expression profiling of RAGP neurons, we identified neural subpopulations, membrane receptors and ion channels that may be targeted by pharmacologic agents upstream of the myocardium. In rat and canine intrinsic cardiac neurons, both nicotinic and muscarinic synaptic transmission has been studied (42, 43). Here, we show that muscarinic receptors are expressed in the RAGP. Hence, when muscarinic antagonists such as atropine are applied, they may not simply affect muscarinic receptors at the atrial myocardium but also in the cardiac neurons. This may provide another explanation for seemingly paradoxical phenomena such as atropine-induce bradycardia (44, 45), as prior studies on the RAGP using atropine to mitigate cholinergic inputs to the myocardium may have inadvertently impacted RAGP neurophysiology (33, 46). Genetic association studies that implicate muscarinic receptors in HR recovery in exercise or ion channels with sudden unexplained death in epilepsy are agnostic to tissue type (47, 48). These variants may serve as markers for pathology in the nervous system rather than the myocardium.

Similarly, the presence of HCN4 channels in cardiac neurons may suggest that the *I*_*f*_ blocker ivabradine may exert its action via *I*_*H*_ in the ICNS in addition to affecting the *I*_*f*_ in the SAN and warrants further study of how HCN4-positive cells in the SAN region influence its pacemaker function (49). Delineation of neuronal subpopulations and molecular states by gene expression profiling suggests phenotypic diversity beyond what is ascertainable by immunohistochemistry or the application of pharmacologic agents (50). Characterizing the plasticity of RAGP arising due to dynamic shifts in these neuronal states may elucidate molecular mechanisms driving pathophysiological response in heart disease to inform novel neuromodulatory interventions.

The present study further advances our knowledge of the functional innervation of the cardiac conduction system and working myocardium. VNS causes a shift in the pacemaker site and activation pattern within the SAN region (51–53). Following the initial description that stimulation at an intercaval region of the RA conveys vagal input to the SAN (4), electrical stimulation of the RAGP fat pad was shown to reduce HR in dogs and humans (6, 54). McGuirt *et al.* showed that while ablation of the RAGP abolishes response to cervical bilateral VNS, VNS was still able to reduce the response to sympathetic stimulation, indicating that RAGP ablation eliminated direct, but not indirect, parasympathetic control of the SAN (55). This highlights the complex interplay between the cardiac GPs and higher levels of the cardiac neuraxis rather than cardiac GPs acting as relay stations (3). A laterality has been ascribed to both vagal and sympathetic inputs to the SAN wherein the right vagus and stellate ganglion influence the SAN, the left vagus predominantly impacts the AVN and the left stellate affects LV inotropy (56, 57), although one study showed left vagal inputs via the RAGP (58). We demonstrated that RAGP ablation impacts bilateral vagal inputs as well as the effect of the right vagus nerve on the AVN. This builds on a prior study in guinea pig that demonstrated a nerve tract extending from SAN, inferior to the ostium of the coronary sinus and toward the posterior aspect of the AVN, although this has been debated (36, 59). Furthermore, as RAGP ablation also impacted LV contractility, our findings continue to challenge the notion that the GPs can be specifically targeted to independently influence SAN or AVN function, as proposed in animal models and human studies (60–62). Although ablation of the region did not significantly impact SGS-induced tachycardia, neurons within the right stellate ganglia were identified after retrograde labeling which is consistent with right SGS-induced increases in SAN rate. Adjunctive ablation of the posterior atrial GP, which is located between the SVC and aorta, may have abolished SGS-induced tachycardia, as murine and canine models have suggested TH-positive fibers course via the medial aspect of the SVC to the SAN (63, 64). As some cardiothoracic surgeons are exploring SAN denervation for SAN ablation for inappropriate sinus tachycardia, these findings suggest that the region of the RAGP-SAN should be spared (65).

We showed that RAGP stimulation typically induces reductions in atrial rate, but we also demonstrated changes in AVN conduction and induction of AF. Interestingly, bradycardia was easier to induce with decentralization following bilateral cervical vagotomies and vagosympathetic trunk debranching (i.e. bilateral stellate ganglionectomy). In the setting of vagotomies only, RAGP stimulation variably induced a mild bradycardia, while decentralization resulted in profound sinus bradycardia with ventricular asystole or AF. This suggests central input impacting the threshold to electrical stimulation and contrasts with prior studies where decentralization reduces response to pharmacologic stimulation (40, 66). Clinically, high frequency stimulation using an endovascular approach to elicit a vagal response can be challenging, which is why some clinical trials have espoused an anatomic-guided approach to GP ablation (61, 62, 67). Here, we show that central input to the ICNS tamps down the ability to induce a response to high frequency stimulation. We further demonstrate this in our human atrial wedge preparation, a ‘decentralized’ state, in which we elicited a response with high frequency stimulation. The response was tachycardic rather than bradycardic and may have been due to increased frequency of stimulation, as a prior study in dogs demonstrated ectopy of increased severity with uptitration of stimulation frequency (68) and as we demonstrated in our porcine model (**Fig S7A**). Alternatively, electrical stimulation may be impacting adrenergic fibers of passage *en passant* to the SAN. For example, studies in humans (i.e. in the absence of decentralization) suggested that RAGP stimulation affects the SAN without AH prolongation (6, 69), while other canine studies describe bradycardia, tachycardia, AH prolongation and atrial fibrillation induced by RAGP stimulation (60, 70, 71). This further highlights the difficulty with specifically targeting only RAGP neurons as opposed to nerve tracts. However, given anatomy-guided identification and ablation of this region in patients, this is a clinically relevant approach. While intramural autonomic nerves in the vicinity of the SAN have been stimulated in an atrial wedge preparation in dog and rabbit to elicit a change in atrial rate, to our knowledge, this is the first description of how the RAGP exerts control over the pacemaker site (72, 73). Importantly, clinical studies evaluating RAGP ablation for sinus node dysfunction have not reported sinus node electrograms to demonstrates effects on the sinus node itself (74).

The development of therapeutics that target the cardiac autonomic nervous system could provide an effective strategy for treating arrhythmias. However, as outlined by the present study, neuromodulatory therapies that target the RAGP to affect the SAN may result in off-target effects. In the AFACT study, in which patients were randomized to AF ablation with or without GP ablation, 5% of patients required pacemaker implantation and two-year follow-up did not show a reduction in AF recurrence (10, 75). Heterogeneity in targets for intervention across studies has yielded variable results, and the approach of ablating GPs does not selectively target neurons, as afferent, efferent and local circuit neurons are likely impacted (9, 76). Inappropriate sinus tachycardia has emerged as a consequence of disrupting parasympathetic fibers to the SAN, although SAN denervation has also been applied to cases of inappropriate sinus tachycardia (65, 77). In studies of GP ablation for sinus bradycardia and syncope, radiofrequency ablation of the RAGP resulted in increased HRs and a 95% reduction in syncope, respectively, but the long-term consequences of such cardioneuroablation are, as yet, unclear (61, 62). Phenotyping of neurons within the RAGP may allow for improved precision in targeting neuronal subpopulations with the goal of addressing specific aspects of cardiac control. One could speculate however that the redundancy of a distributed network of GPs maintains cardiac homeostasis.

This study affirmed the powerful cholinergic control of the RAGP over the SAN, and, for the first time, provides a roadmap of gene and protein expression profiles as well as functional assays of RAGP effects on whole heart and SAN function. The work identifies the RAGP as an integrative neural structure as opposed to relay station within the distributed network of the ICNS. This study provides a foundation upon which further exploration of the functional implications of the diverse neural population influences cardiac physiology and disease. This approach provides a template that may be adopted for defining other components of the ICNS to identify therapeutic targets within the autonomic nervous system to ameliorate cardiovascular disease.

## Acknowledgements

We would like to thank Sarah Hiyari, PhD and Amiksha Gandhi for their contributions to the tissue clearing of porcine and human specimens, respectively, and Toru Adachi, MD; Taro Temma, MD and Christopher Chan for their assistance with the porcine experiments. We extend special thanks to Warwick Peacock, MD; Michael C. Fishbein, MD; and the UCLA Donated Body Program for assistance with autopsy studies. We are also appreciative of the assistance of the California Nanosystems Institute for microCT imaging and the UCLA Broad Stem Cell Research Center Microscopy Core for imaging analysis. We thank James Weiss, MD; Noel Boyle, MD, PhD; and Olujimi Ajijola, MD, PhD for their comments on the manuscript.

