## Supplemental Tables and Figures for "Innervation and Neuronal Control of the Mammalian Sinoatrial Node: A Comprehensive Atlas"

**Supplemental Material**

| **Table S1. Primary Antibodies** | | | | | | |
| --- | --- | --- | --- | --- | --- | --- |
| Antibody | Host | Immunogen | Company | Catalog No. | RRID | Dilution |
| Anti-Vesicular Acetylcholine Transporter (VAChT) | rabbit | aa 475-530 from rat VAChT | Synaptic Systems | 139 103 | RRID: AB_887864 | 1:500 |
| Anti-Tyrosine Hydroxylase (TH) | sheep | Native TH from rat pheocromocytoma | Millipore | AB1542 | RRID: AB_90755 | 1:500 |
|  | rabbit | SDS-denatured rat tyrosine hydroxylase, purified from pheochromocytoma | Pel-Freez Biologicals | P40101-150 | RRID:AB_2313713 | 1:1000 |
| Anti-Vesicular Monoamine Transporter 2 (VMAT2) | rabbit | Synthetic peptide (aa 1-20 from mouse VMAT2) | Synaptic Systems | 138313 | RRID: AB_2619826 | 1:200 |
| Anti-Neuropeptide Y (NPY) | rabbit | Synthetic NPY coupled to bovine thyroglobulin (BTg) | ImmunoStar | 22940 | RRID: AB_2307354 | 1:1000 |
| Anti-Microtubule Associate Protein 2 (MAP2) | chicken | Recombinant fragment of human MAP2, aa 235-1588 | Abcam | ab5392 | RRID: AB_2138153 | 1:1000 |
| Anti-Substance P (SP) | rabbit | Synthetic SP coupled to KLH | ImmunoStar | 20064 | RRID: AB_572266 | 1:1000 |
| Anti-Vasoactive Intestinal Peptide (VIP) | rabbit | Porcine VIP coupled to BTg | ImmunoStar | 20077 | RRID: AB_2572270 | 1:1000 |
| Anti-Somatostatin | rabbit | Synthetic SOM coupled KLH | ImmunoStar | 20067 | RRID: AB_572264 | 1:1000 |
| Anti-Neuronal Nitric Oxide Synthase (nNOS) | goat | aa 1423-1434 of human NOS1 | Abcam | ab1376 | RRID: AB_300614 | 1:1000 |
| Anti-Protein Gene Product 9.5 (PGP9.5) | rabbit | Synthetic peptide to residues in Human PGP9.5 | Abcam | ab108986 | RRID:AB_10891773 | 1:500 |
| Anti-S100 | rabbit | S100 isolated from cow brain | Dako | GA504 | RRID:AB_2811056 | 1:400 |
| Anti-Choline Acetyltransferase (chAT) | goat | Human placental enzyme | Millipore | AB144P | RRID:AB_2079751 | 1:25 |
| Anti-Calcitonin Gene-Related Peptide (CGRP) | goat | Rat CGRP C-terminal peptide,  VKDNFVPTNVGSEAF | Abcam | ab36001 | RRID: AB_725807 | 1:1000 |
|  | mouse | Rat alpha-CGRP | Abcam | ab81887 | RRID: AB_1658411 | 1:1000 |

**Table S2. Primers for neuron-specific genes used in high-throughput real-time PCR experiments**

| **Gene** | **Forward Primer** | **Reverse Primer** |
| --- | --- | --- |
| *NeuN* | GTTTACCTCCCAACCCGAGG | CTCGGCTGTGCCACTTTTTC |
| *PGP9.5* | TGCCTTTTCCGGTGAACCAT | AACGGGGATAAAGCGAAGGG |
| *MAP2* | TGCCGGGGAAGGTGTACTTA | TCTTCTTCTGCGCCTTGCAT |

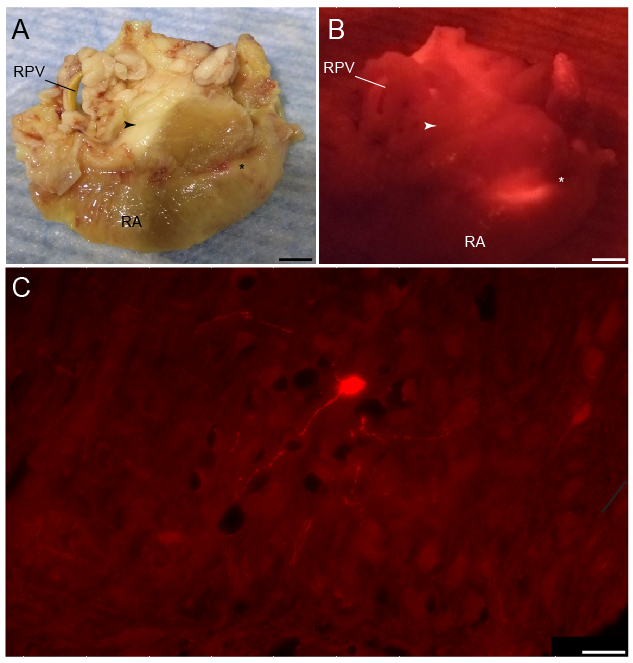

**Fig. S1** **Injection site of DiI retrograde tracer in the SAN region.** **A** Brightfield photomicrograph of anterior view of heart showing RAGP (arrowhead) and SAN (*) region. **B** Fluorescence photograph of the same regions (RAGP: arrowhead; SAN: *) showing the DiI injection site. SVC, superior vena cava; RA, right atrium. **C** Confocal image of a neuron in right stellate ganglion labeled with DiI. Scale bar are 5mm (**A, B**) and 200µm (**C**).

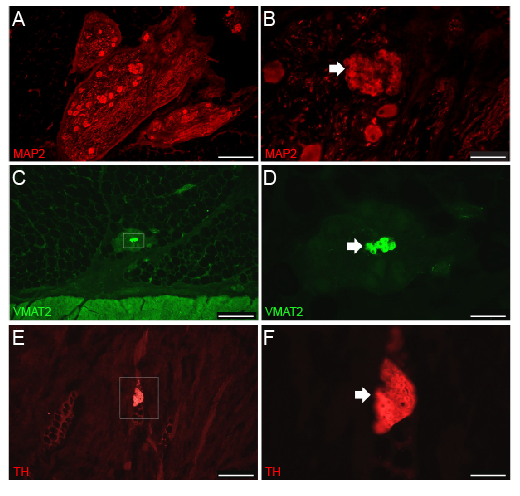

**Fig. S2 Small intensely fluorescent (SIF) cells were identified based on labeling for noradrenergic markers and MAP2, their small size and frequent clustering.** SIF cells were found at various sites throughout RAGP sections: between neurons in the ganglia, in the atrial muscle and amongst fat cells. **A** Low magnification fluorescence microscopic image of ganglia labeled for MAP2. **B** Cluster of SIF cells shown at higher magnification in panel B. SIF cells (arrow) are much smaller than neurons in the same field. **C** Low-magnification fluorescence microscopic image of small ganglion with VMAT2-positive SIF cells. **D** SIF cell cluster stained for VMAT2 (arrow) at higher magnification. **E** Cluster of TH-positive SIF cells located between atrial muscle fibers. **F** TH-positive SIF cell cluster (arrow) shown at higher magnification.

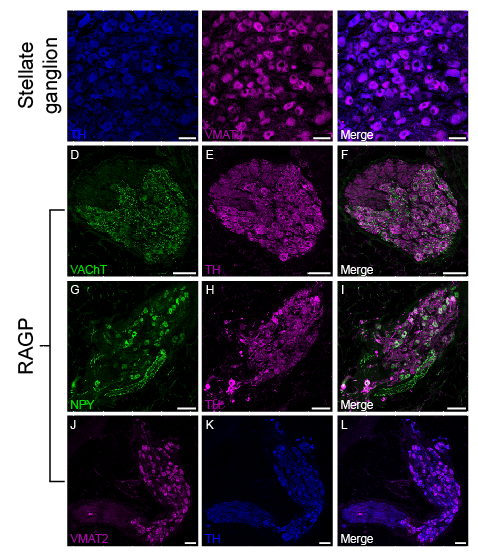

**Fig. S3 Noradrenergic neurons in the stellate ganglia and atypical RAGP ganglion that contained noradrenergic neurons.** **A-C** Confocal images of double labeled section from porcine stellate ganglion showing that TH and VMAT2 are colocalized in noradrenergic neurons. **D-L** Atypical RAGP ganglion that contained noradrenergic neurons. **D-F** Confocal images of ganglion double labeled for VAChT (**D**) and TH (**E**) shows strong labeling of all neurons for TH. (**F**) Overlay image shows cholinergic varicosities around TH-positive neurons. **G-I** Confocal images of the same ganglion double labeled for NPY (**G**) and TH (**H**) shows many neurons labeled for each marker. (**I**) Merged image shows colocalization of NPY and TH in cell bodies. **J-L** Confocal images of same ganglion double labeled for VMAT2 (**J**) and TH (**K**). **L** Merged image shows extensive colocalization of both markers in this ganglion. Scale bars are 50µm (**A-C**) and 100µm (**D-L**).

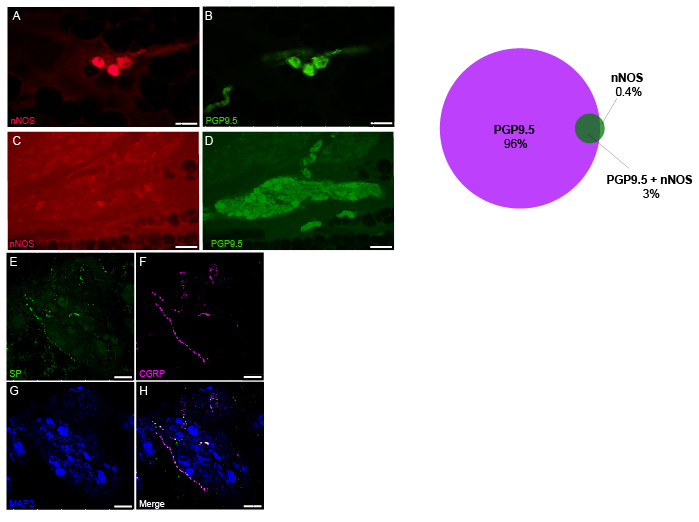

**Fig. S4** **nNOS is found in only a few neurons in the porcine RAGP and CGRP and SP are found in nerve fibers.** **A-D** Fluorescence microscopic images of sections double labeled for nNOS and PGP 9.5. **A, B** Neurons that stain strongly for nNOS. **C, D** Images from a typical ganglion that contains several PGP 9.5-positive neurons that lack nNOS. **E** Venn diagram illustrates proportion of PGP9.5-positive, nNOS-positive and both PGP9.5- and nNOS-positive neurons **F-I** CGRP and SP are colocalized to varicose nerve fibers in porcine RAGP, but these do not surround ganglionic neurons. Confocal images of a ganglion that was triple labeled for SP, CGRP and MAP2. **F-G** Varicose nerve fibers staining for SP and CGRP, respectively. **H** Staining for MAP2 shows neuronal cell bodies and their processes. **I** Merged image shows the colocalization of SP and CGRP to nerve fibers and the close apposition of some nerve fibers to MAP2-positive neurons and processes. Neither SP nor CGRP is colocalized with MAP2. Scale bars are 50µm (**A-B, F-I**) and 100µm (**C-D**).

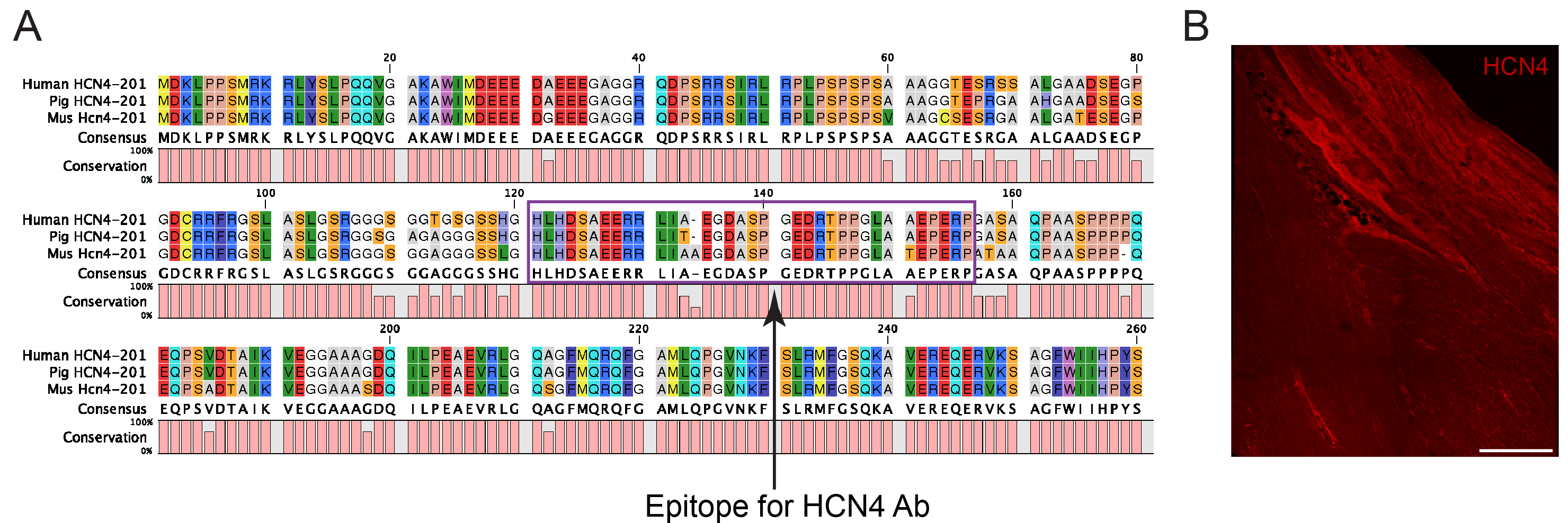

**Fig. S5 Immunohistochemical assessment of porcine SAN.** **A** Sequence homology comparing the *HCN4* gene in human, pig and mouse. Note that the human epitope used to generate the HCN4 antibody (Alomone labs, APC-052) is very similar to that of pig and mouse. **B** Immunostaining of HCN4 in porcine SAN shows poor contrast. Scale bar: 500µm.

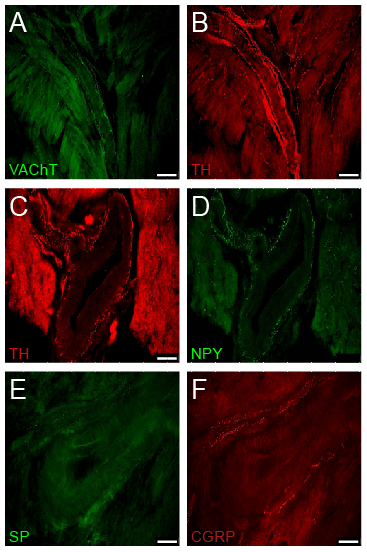

**Fig. S6 A-D** Atrial blood vessels are innervated by cholinergic, noradrenergic, and peptidergic nerve fibers. **A, B** Double labeling for VAChT and TH shows cholinergic and noradrenergic nerves around vessel cut longitudinally. **C, D** Double labeling for TH and NPY shows that these markers are colocalized around a blood vessel cut in cross section. **E, F** Double labeling for SP and CGRP shows that both sensory neuropeptides are present around a vessel cut in cross section, but CGRP is more abundant. Scale bars are 50µm (**A-B**) and 100µm (**C-F**).

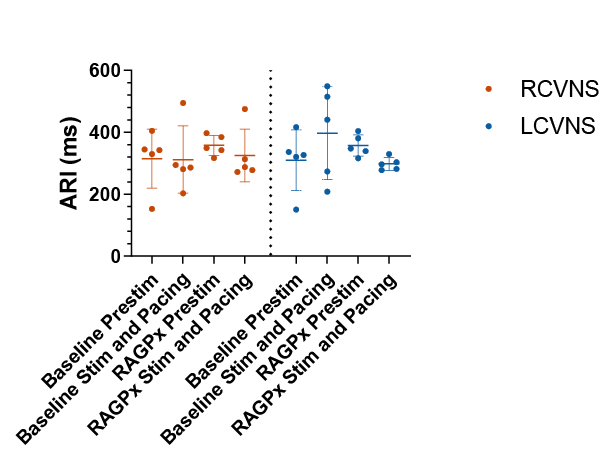
**Fig. S7 RAGP ablation did not impact VNS-induced changes in ventricular ARI during atrial pacing**

No significant changes in average global ventricular ARI before and after RAGP ablation and before and during either RCVNS or LCVNS with concurrent atrial pacing were identified. Prestim: prior to VNS, RAGPx: RAGP ablation.

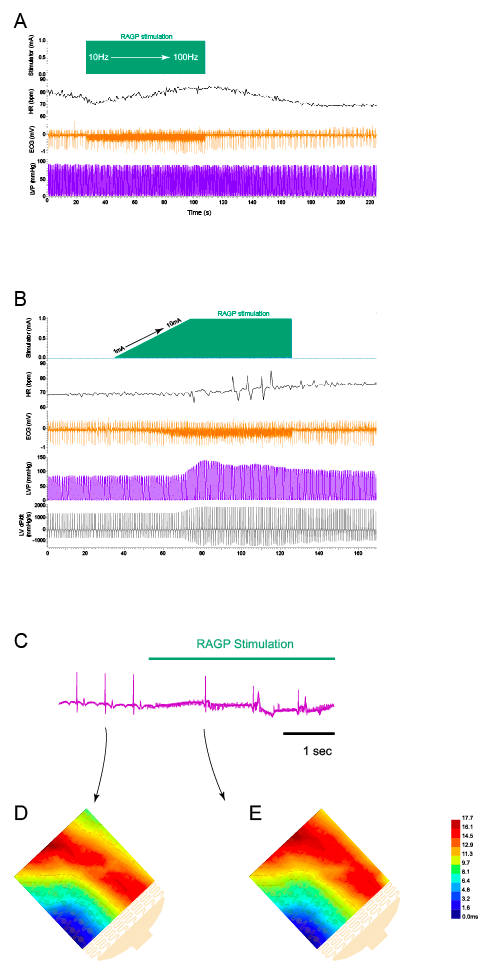

**Fig. S7 Stimulation of RAGP elicits different responses from the SAN and LV**

**A** Example of bradycardia elicited by lower frequency (10Hz) RAGP stimulation with tachycardia at higher frequency (100Hz) stimulation. **B** Increase in current from 1 to 10mA with mild increase in HR and significant increase in LV pressure and contractility (dP/dt). **C** Example of RAGP stimulation-induced bradycardia. Activation map before (**D**) and during (**E**) RAGP stimulation.

**Supplemental Movie Legends**

**Movie S1 Network of ganglia and interconnecting nerve fibers are found in the porcine RAGP.** 3D projection of portion of iDISCO+-cleared porcine RAGP immunostained with PGP9.5 (purple).

**Movie S2 3D projection of ganglion found in porcine RAGP**. iDISCO+-cleared ganglion immunostained with PGP9.5 (purple) at higher magnification.

**Movie S3 Ganglia in the human RAGP are found within the adipose tissue and the fat-muscle interface.** 3D stack of sequential hematoxylin and eosin sections of human RAGP demonstrating ganglia embedded in fat as well as at the fat-posterior right atrial wall interface.

**Movie S4 Network of ganglia and interconnecting nerve fibers are found in the human RAGP.** Portion of iDISCO+-cleared human RAGP immunostained with PGP9.5 (purple).

**Movie S5 The sinoatrial nodal artery (SNA) supplies the human RAGP.** MicroCT image sequence of contrast-injected right coronary artery (RCA; red) with sinoatrial nodal branch supplying the RAGP (yellow) of a human cadaveric cardiac specimen.

**Movie S6 The porcine SAN is densely innervated.** 3D projection of portion of porcine SAN immunostained with PGP9.5 (yellow). Muscle autofluorescence is purple. Note the multiple ganglia found in the SAN region. Large nerve fibers can be seen along the crista terminalis. ST: sulcus terminalis, SVC: superior vena cava.
